# Coding of the basic components of subjective value in primate dopamine neurons: subjectively weighted reward amount and probability

**DOI:** 10.1101/2025.02.21.639529

**Authors:** Simone Ferrari-Toniolo, Leo Chi U Seak, Wolfram Schultz

**Affiliations:** Department of Physiology, Development and Neuroscience, University of Cambridge, Cambridge, UK

**Keywords:** Dopamine, reward, choice, utility, weighted probability, heterogeneity, economic axioms

## Abstract

Behavioral choices of uncertain rewards suggest that agents construct subjective reward value by combining the basic value components of utility and weighted probability. Despite the general acceptance of this evaluation mechanism for explaining economic choice, knowledge about its neuronal implementation is fractionated and remains essentially unknown. We investigated in monkeys whether reward signals in dopamine neurons might represent subjective reward value based on these two fundamental value components. Despite some heterogeneity across individual neurons, the dopamine population signal reliably represented the axiomatically defined integration of utility and weighted probability into subjective value in a way that closely matched the animal-specific choice behavior. In particular, we identified a crucial contribution of subjectively weighted probability to the dopamine signal of subjective reward value. These data demonstrate a neuronal implementation of subjective value constructed from the two most basic subjective reward components.

**Preprint Server:** An earlier version of this manuscript has been uploaded to bioRxiv in February 2025 (CC-BY).

## INTRODUCTION

The essence of value-based decision making lies in assigning a unique value to each choice option we face. This evaluation allows us to select the option that holds the greatest value for us. This conceptually simple idea has been formally developed into rigorous economic models. However, the neuronal implementations of these concepts are only incompletely known.

Economic choice models define how reward components are combined into subjective values. Two of the most fundamental value components are reward magnitude (the physical amount of reward) and reward probability (the likelihood of obtaining the reward). Expected utility theory (EUT) (von Neumann and Morgenstern, 1944) represented the first axiomatic formulation of subjective values. According to EUT, reward magnitudes are subjectively modified into utilities, representing a nonlinear transformation of the objective reward quantities. An option’s subjective value is defined as its expected utility (EU), i.e., the utility of its outcome weighted by its probability of occurrence. EUT’s four axioms define the necessary and sufficient conditions supporting this expected utility representation. Given a specific utility function, this elegant formulation of subjective values can fully predict choices between uncertain rewards.

However, choices deviate from EUT predictions in specific situations. In a prominent example (Allais, 1953), human subjects violated the independence axiom of EUT, which states that preferences between two options should not change after adding the same probability of a common outcome to both options. A subjective transformation of the objective reward probabilities was introduced to explain these unexpected choice behaviors. The subjectively weighted probability became a crucial component of prospect theory (Kahneman and Tversky, 1979) and of other non-EU theories (Starmer, 2000). These economic models relied on the subjective weighting of both reward magnitude and probability, representing a more complete account of subjective value in human and animal decision making (Barberis, 2013; Stauffer et al., 2015).

The subjective nature of economic value is manifested in subject-specific risk attitudes, a crucial feature of individual choice behavior in uncertain situations. Economic risk (i.e., the variance in an option’s statistical reward distribution) is higher when outcomes are less predictable. In choices between a sure reward and a risky option with equal objective expected value (EV), preferring the sure reward identifies risk avoidance, while choosing the risky option identifies risk seeking. The latter choice could be driven by overweighting of reward probability as well as by underweighting of sure reward magnitude. In formal economic terms, risk attitudes are determined by the relative shapes of utility and weighted probability functions (Tversky and Kahneman, 1992). Thus, to fully understand the source of risk attitudes, it is crucial to consider the subjective weighting of both reward magnitude and probability.

Neuroeconomic research in the past 25 years has explored the hypothesis of a value-based principle implemented in human and animal brains (Serra, 2021). Midbrain dopamine neurons have emerged as strong candidates for encoding subjective values. Their activations, strongly modulated by unexpected rewards, are compatible with a value-updating role in reinforcement learning (Schultz, Dayan and Montague, 1997; Glimcher, 2011). The responses of primate dopamine neurons reflect utility (Stauffer, Lak and Schultz, 2014). Reinforcement learning models employing nonlinear reward magnitudes can explain empirical risk preferences (Louie, 2022). These results suggest that the dopamine teaching signal may contribute to value-learning, ultimately driving decisions towards subjectively optimal rewards. Yet, this leaves an incomplete picture of the dopamine value signal, as the crucial contribution of subjectively weighted reward probability has not been considered. Only the combined coding of subjective reward magnitude and probability would suggest a neuronal valuation mechanisms compatible with formal economic theories of choice.

Here we investigated whether monkey midbrain dopamine neurons encode the basic components of subjective value, namely utility and weighted probability, combining them into an integrated neuronal signal of subjective reward value. We first compared behavioral and neuronal value measures based on the economic axioms of EUT, which mathematically define subjective values. We then quantified the nonlinear responses to both reward magnitude and probability to evaluate how their subjective values are encoded in dopamine neurons. We found that the dopamine population activity reflected subjective values across options with broadly varying reward magnitudes and probabilities, regardless of their physical composition. The dopamine responses coded both utility and subjectively weighted probability. A neuronal test of the crucial independence axiom of EUT, which is violated when probabilities are not subjectively weighted, confirmed the crucial contribution of nonlinear probability weighting to neuronal value computations: the dopamine signal matched behavioral choices only with subjective nonlinear probability weighting. Together, these results indicate that responses of primate dopamine neurons represent the integration of utility and weighted probability, the two fundamental components of subjective economic value.

## RESULTS

### Design and behavior

This study aimed to advance the existing knowledge about the components of reward value signals in primate dopamine neurons. Using a stringent economic axiomatic approach, we related neuronal activities to behaviorally elicited reward value measures. We used the continuity axiom of EUT to formally define and quantify subjective values assigned to risky choice options (gambles). The continuity axiom (Eq. 1) states that decision makers should be indifferent in choices between one option and a probabilistic combination of two other options, reflecting the notion that subjective values originate from a graded combination of the option’s attributes. The axiom implies the existence of a real-valued subjective value function representing the ranking of choice options (Camerer, 1989; Weber and Camerer 1987). Our preceding behavioral study (Ferrari-Toniolo et al., 2021) demonstrated compliance of monkey’s choices with the continuity axiom.

Following the axiom definition, we estimated subjective values from choices for a broad set of options. Applying the same principle to neurons, we identified a corresponding neuronal subjective-value measure. These behavioral and neuronal foundations were the basis for directly relating neuronal responses to choice behaviors.

In each trial, the monkey chose between two choice options, which corresponded to specific combinations of reward magnitude (m, 0 – 0.5 ml) and reward probability (p, 0 – 1). The two options were presented at pseudorandomly alternating fixed left and right positions (Figure 1A, B). One option was a sure reward with magnitude m (safe option; p = 1). The alternative option was a probabilistic reward (gamble, p < 1) with two possible outcomes: either reward magnitude m with probability p, or no reward with probability 1 – p.

**Figure 1.**
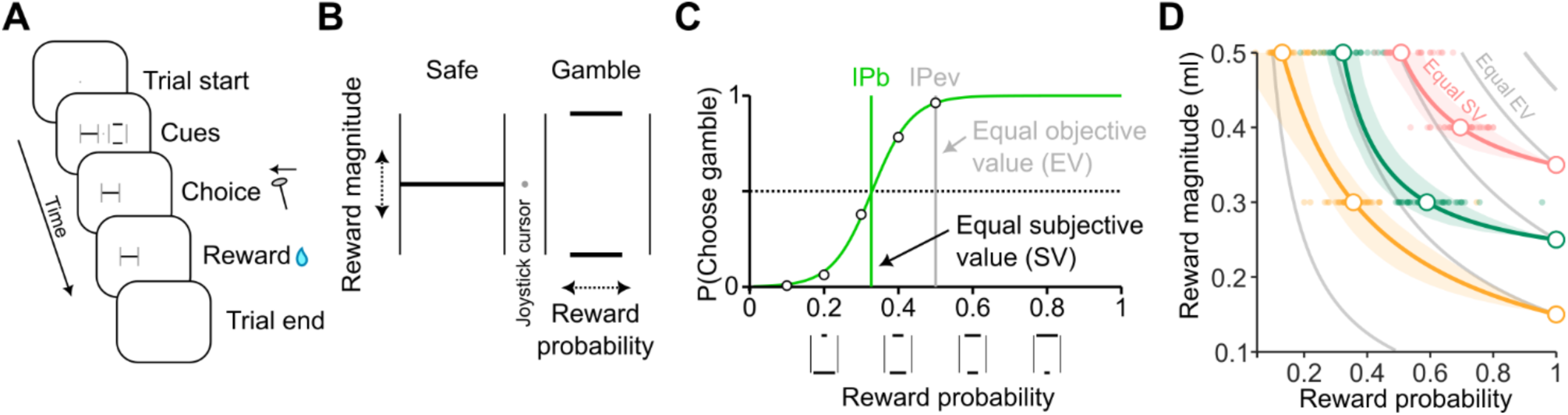
Experimental design and choice-indifference data. A) Trial structure. The animal was trained to control a cursor on the screen through left-right joystick movements. After holding the cursor in a central position for a variable time (1 to 1.5 s), two visual cues, indicating the two choice options, appeared. The preferred option was indicated by moving the joystick to the corresponding side, while the unchosen option disappeared. After a fixed delay (1 s) the reward corresponding to the chosen option was delivered, followed by a further blank-screen delay (1 s) and by a variable intertrial period (1 – 2 s). B) Visual cues. Each bar represented a possible outcome magnitude (vertical position; 0 – 0.5 ml) and probability (horizontal width). A single bar indicated a sure reward (safe option), two bars corresponded to a probabilistic reward (gamble option). C) Indifference point estimation. In choices between a fixed safe option and a gamble with varying reward probability and fixed magnitude, we estimated the behavioral indifference point (IPb) as the probability for which the two options were equally chosen, i.e., had the same subjective value (SV). IPev: probability for which the two options had the same objective expected value (EV). D) Magnitude-probability indifference curves (ICs). IPb’s relative to different gamble magnitudes (dot: single session IPb; circle: IPb from all sessions, N = 47) resulted in a series of equal-SV points. Fitting a parametric function (Eq. 3) identified the corresponding IC. Varying the safe option identified different ICs (colored curves), which significantly deviated from the objective equal-EV lines (areas: bootstrap 95% CI, N = 1,000 resamples with replacement). Data from Monkey A. See Figure S1 for Monkey B data.

We used observable behavioral choice to infer subjective values in line with the continuity axiom principle. In choices between a safe and a gamble option, we defined the subjective economic value of the safe option as the gamble’s reward probability for which the two options were chosen equally frequently (P(choice) = 0.5). For different gamble probabilities we computed the probability of choosing the gamble option (P(choose gamble)) and fitted a softmax function (Eq. 2) to estimate the gamble probability at the behavioral indifference point (IPb) (Figure 1C). The estimated IPb represented the numeric subjective economic value of the safe reward in relation to the gamble outcomes. Importantly, when preferring the gamble with physical lower mean reward (expected value, EV) animals forwent the objectively larger safe reward, demonstrating reliance on subjective value rather than physical value. This also indicated a risk seeking attitude, resulting in an IPb lower than the corresponding IPev (i.e., the gamble probability for which the gamble EV matched the safe magnitude) (Figure 1C). We used this IPb measure, quantifying the animal’s subjective evaluation of choice options, for the quantitative comparison between behavioral and neuronal data.

We defined a two-dimensional magnitude-probability space, in which each point corresponds to a gamble. The full set of IPb’s identified an indifference map, i.e., a series of indifference curves (ICs), connecting gambles with equal subjective value in this magnitude-probability space (Figure 1D; colored lines). If the animal only considered either magnitudes or probabilities when making choices, the ICs would be linear and parallel to the respective axis; any other pattern would instead indicate the integration of the two reward components into a subjective value. The distance between equal subjective value curves (i.e., ICs) and equal-EV curves highlighted the subjective nature of economic values underlying choices. We found ICs leaning towards the left side of equal-EV curves (Figure 1D), thus identifying a general risk-seeking attitude.

Overall, these data show that monkeys made meaningful choices which reflected a subjective integration of objective reward magnitude and probability information. The choice-indifference method, based on the continuity axiom principle, numerically quantified subjective reward values, crucial for relating these subjective quantities to the corresponding neuronal measures.

### Indifference points as neuronal subjective value measures

Dopamine neurons in the primate midbrain respond to temporally unpredicted reward-predicting cues, consistent with their coding of a temporal reward prediction error (Schultz et al., 1997; Schultz, 2024). To test whether dopamine cue responses are compatible with economic-value coding, we defined a neuronal subjective value measure, the neuronal indifference point, which we then compared with the corresponding behavioral measure. These two measures were based on the same economic principle, the continuity axiom of EUT, for a coherent and direct comparison between neuronal and behavioral quantities.

We recorded the activity of 172 dopamine neurons from two macaque monkeys (N = 59 in monkey A, N = 113 in monkey B), while presenting either two reward-predicting cues (choice task, described earlier) or a single reward-predicting cue (single-option task, see Methods). Individual dopamine neurons were identified from their anatomical location, low baseline activity rate, positive response to unexpected rewards and elongated waveforms (see Methods). As expected, the cue response of a typical dopamine neuron increased with both reward magnitude and probability, with peak response around 150 ms after cue onset (Figure 2A). For all data analyses, we selected neurons with significant multiple-linear regression coefficients for reward magnitude and probability at cue presentation (t test on m and p beta coefficients, P < 0.05; see Methods), separately tested in single-option and choice tasks (single-option task: N = 54 (out of 56) in monkey A, N = 48 (out of 59) in monkey B; choice task: N = 51 (out of 59) in monkey A, N = 56 (out of 66) in monkey B).

**Figure 2.**
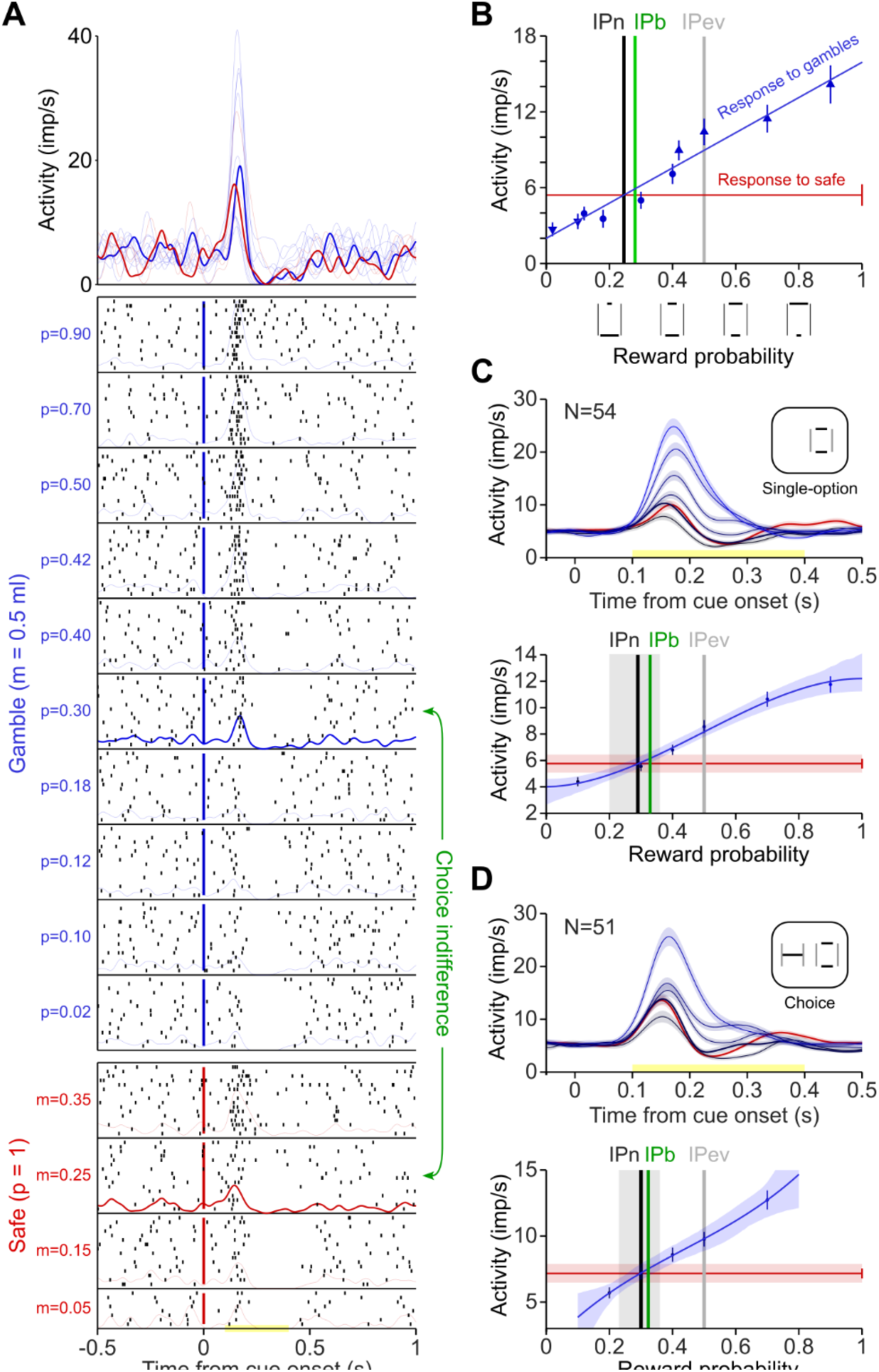
Elicitation of behavioral and neuronal indifference points. A) Example dopamine neuron response. Raster plot of the action potentials of an individual dopamine neuron in response to a visual cue representing either a safe option (red) or a gamble option (blue), during single-option trials. Colored curves: gaussian-smoothed spike density functions. Larger safe reward magnitudes and gamble probabilities produced an increased neuronal response, in line with the expected coding of reward prediction error in dopamine neurons. ‘Choice indifference’ arrows (green) link two options that were equally chosen by the monkey, implying same subjective value. A neuron coding subjective value is expected to be equally active for two equally-chosen options. B) Neuronal indifference points. The neuronal indifference point (IPn) was defined as the gamble probability for which the neuronal response to the safe option (m = 0.25 ml) matched the response to the gamble option (m = 0.5 ml). The IPn was computed as the intersection between two lines: the regression line of the neuronal activity for different gamble probabilities (blue) and the line representing the mean response to the safe option (red). In this example neuron (same neuron as in panel A) the IPn closely matched the behavioral indifference point (IPb). Behavioral (and neuronal) indifference points were lower than the objective IPev, revealing (and coding) a risk-seeking attitude. C, D) Population activity and IPn. Top: average activity (area: SE) across the neuronal population in response to one safe option (red) and different gamble probabilities (blue). Bottom: estimation of IPn from neuronal population-averaged activity, following the procedure described in panel B. Data from Monkey A in single-option task (C) and choice task (D). In both tasks the population IPn matched the IPb (grey area: bootstrap 95% CI, N = 1,000 resamples with replacement) while significantly deviating from the objective IPev.

To test whether neuronal responses were compatible with the coding of subjective values, as opposed to objective values or values incompatible with revealed preferences, we compared the response to a specific safe option with the response to gambles with different probabilities. The response of a neuron encoding subjective value was expected to be similar between two options for which the subject expressed choice indifference. Indeed, in an example neuron, we observed similar response profiles for the safe option (m = 0.25 ml) and for the gamble (m = 0.5, p = 0.3) among which the monkey was behaviorally indifferent (Figure 2A). To test this more formally, we linearly fitted the responses to different reward probabilities, and calculated the intersection between the fitted line and the safe-option response (Figure 2B). This identified our neuronal value measure, the neuronal indifference point (IPn), which, in subjective-value coding neurons, should match the behavioral indifference point (IPb). In our example neuron, the IPn was close to the IPb, while they both differed substantially from the IPev (Figure 2B). The example neuron thus closely represented the subjective value revealed by the monkey choices, which differed significantly from the objective reward values.

At a neuronal population level, averaging responses across neurons confirmed the match between neuronal and behavioral indifference points (IPb within the 95% confidence interval of bootstrapped IPb, see Methods) during both the single-option task (Figure 2C) and the choice task (Figure 2D). For the choice task, we used as analysis variables the magnitude and probability of the option chosen in each trial. Thus, during choice, dopamine neurons encoded the subjective value of the chosen option. Note that the significant difference between IPn and IPev confirmed that the risk-seeking attitude expressed by the monkey was reflected in the dopamine neurons’ activity.

To test for robustness in the match between our behavioral and neuronal indifference point measure, we repeated the neuronal IPn measurements for different sets of safe and gamble magnitudes and analyzed their correlation with the corresponding behavioral IPb’s (Figure 3A, B). We found a significant correlation between neuronal and behavioral indifference points in both single-option task (Pearson’s rho = 0.995, P = 5.6e-5) and choices (rho = 0.985, P = 3.4e-4).

**Figure 3.**
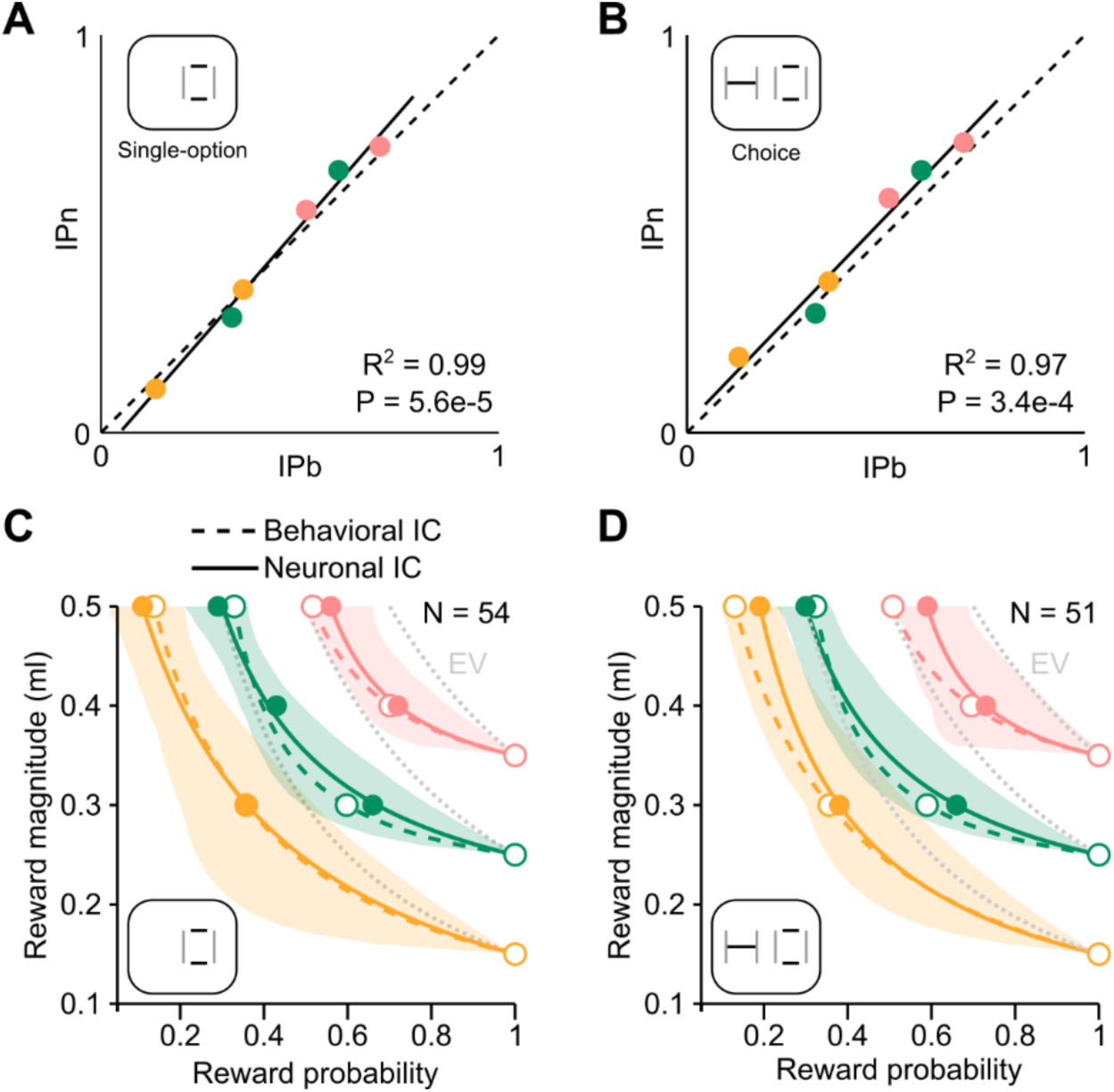
Match between neuronal and behavioral indifference points. A, B) Correlation between neuronal and behavioral subjective value measures. Subjective values (indifference points) were elicited from choice behavior (IPb) and from the averaged neuronal population activity (IPn) during single-option task (panel A) and choice task (panel B). Dashed line: diagonal representing perfect neuronal-behavioral match. Continuous line: linear regression. C, D) Comparison of neuronal and behavioral indifference curves (ICs). Behavioral and neuronal ICs computed with the same procedure by fitting a parametric function (hyperbola) to the indifference points. Area: 95% confidence intervals (CIs) for neuronal ICs. Neuronal ICs matched the behavioral ICs (within the 95% CIs), while significantly differing from the equal expected value (EV) curves (dotted). Data from monkey A. See Figure S2 for monkey B data.

We represented the IPn’s within the same magnitude-probability space which we defined for IPb’s (see Figure 1D) and fitted a series of neuronal ICs to the IPn’s. The neuronal ICs were close to the behavioral ICs, which lay within the confidence interval of the neuronal curves in both the single-option and choice tasks (Figure 3C, D). Thus, the neuronal subjective value measures (IPn) matched the behavioral ones (IPb) across the full range of tested reward magnitudes and probabilities.

To test for the validity of neuronal subjective value coding, we compared the responses of dopamine neurons across different points of the magnitude-probability space, in relation to the behaviorally elicited indifference curves. In particular, we tested whether the neuronal responses were compatible with subjective value differences across ICs, while controlling for objective value differences. The subjective value is constant along each indifference curve and becomes higher for curves closer to the top right of the graph.

Correspondingly, neuronal activities representing subjective values should not vary along ICs and should increase between lower and higher ICs, regardless of the physical composition of the gambles. We tested this hypothesis by statistically comparing (t test, P < 0.05) the average dopamine population response between pairs of gambles lying on either two ICs or the same indifference curve. We found stronger responses for gambles with higher m and p (i.e., higher physical and subjective value) (Figure 4A), higher subjective value despite partly lower physical value (m or p) (Figure 4A_1_), higher subjective value despite equal objective value (EV) (Figure 4A_2_), and similar responses (non-significant difference) to options with similar subjective value despite different EV (Figure 4A_3_).

**Figure 4.**
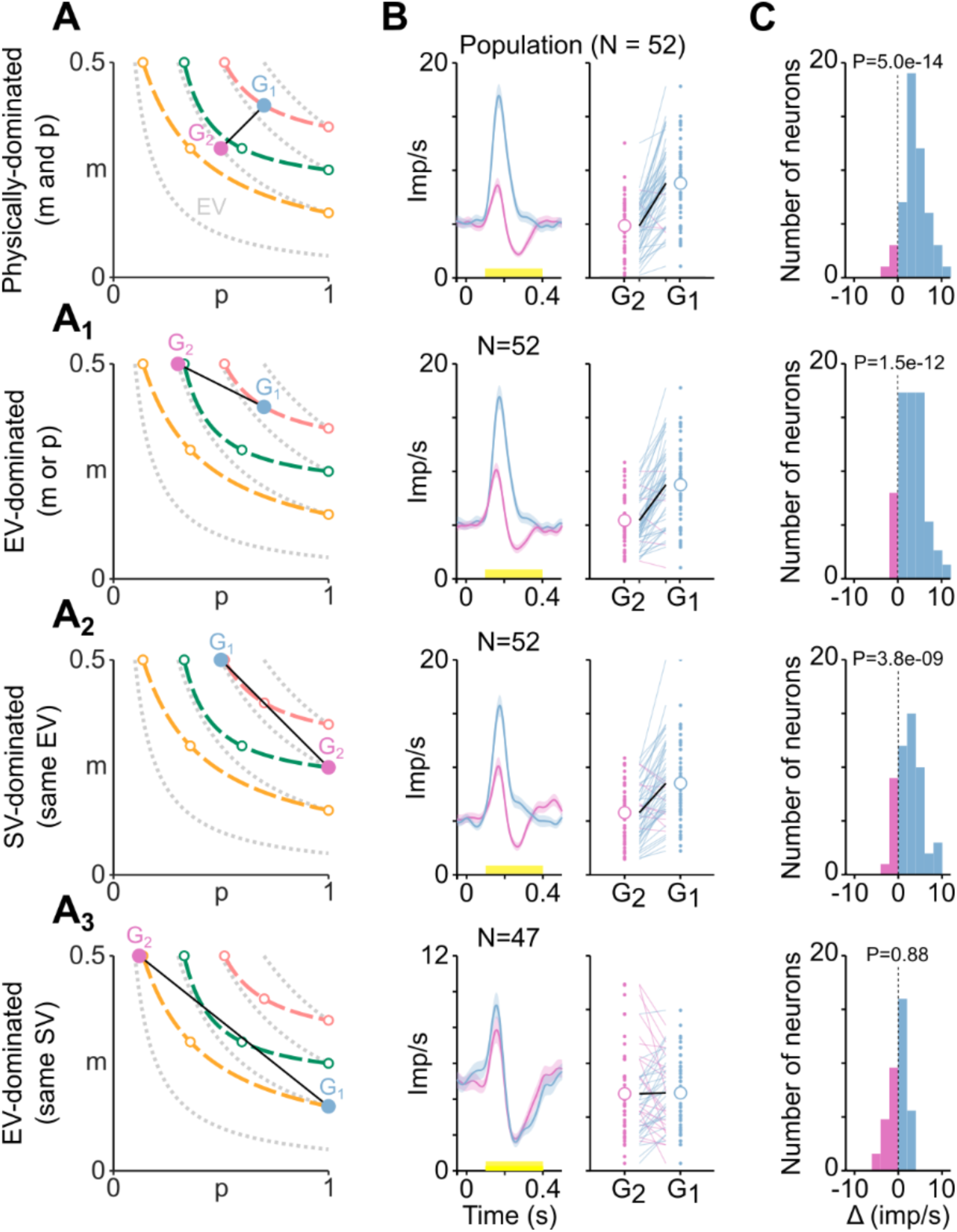
Neuronal responses reflect subjective value rather than physical value. A) Representation of two gambles (G1, G2) in the magnitude-probability space, in relation to the behavioral indifference curves (dashed) and the equal-EV curves (gray). G1 ‘dominates’ G2 (i.e., it has larger objective or subjective value compared with G2) in terms of: A) objective magnitude and probability, A1) EV with smaller magnitude, A2) subjective value (SV) (i.e., indifference curve) with same EV, A3) EV with same SV. B) Neuronal responses to G1 and G2. The average neuronal population response (left) and single neurons’ responses (right) reflect the subjective ranking of the two gambles, with higher activity for the most preferred option. Yellow line (left): time window for computing each neuron’s activity rate (dots, right). Lines (right) connect responses to G1 and G2 for each neuron (color: option for which the neuron showed a stronger response). Black line connects population-averaged activities for the two options (circles). C) Statistical comparison of responses to G1 and G2. Histogram of the number of neurons responding more strongly to G1 (blue) or to G2 (pink). P, statistical p-value of t test on G1 – G2 activity difference (ι1). Population activity was significantly higher for the preferred option. At choice indifference the activity was not significantly different, despite G1 and G2 having different objective value (EV). Data from monkey A during single-option task. See Figure S3 for data from monkey B.

Taken together, these results indicate that dopamine neurons encoded the subjective values of choice options as revealed by choice, regardless of their physical composition in terms of objective reward components.

### A neuronal value code from subjective reward components

The subjective weighting of reward magnitude and probability is a crucial feature of several modern economic choice models. The integration of these subjective reward components define subjective values that explain individual preferences between risky options, suggesting a possible source for individual risk attitudes. To test whether dopamine neuron’s signals reflected these subjectively weighted reward components, we analyzed the nonlinearity in the neuronal responses to different levels or reward magnitudes and probabilities. We then constructed a neuronal indifference map using these nonlinear responses, and compared it with the behaviorally inferred indifference map.

We fitted cubic spline functions, avoiding arbitrary parametric assumptions, to the neuronal population responses. We found generally s-shaped response profiles for both magnitudes and probabilities (Figure 5A, B), with an initial concave portion, followed by linear and then convex regions. Similar profiles were found for single-option and choice tasks. Neuronal responses differed marginally between animals, with monkey B’s neurons, compared with monkey A’s, showing a more pronounced flat response to low probabilities, and a more convex shape in the high magnitude range. We used a bootstrap procedure (resampling with replacement across individual neurons) to account for the variability in neuronal responses.

**Figure 5.**
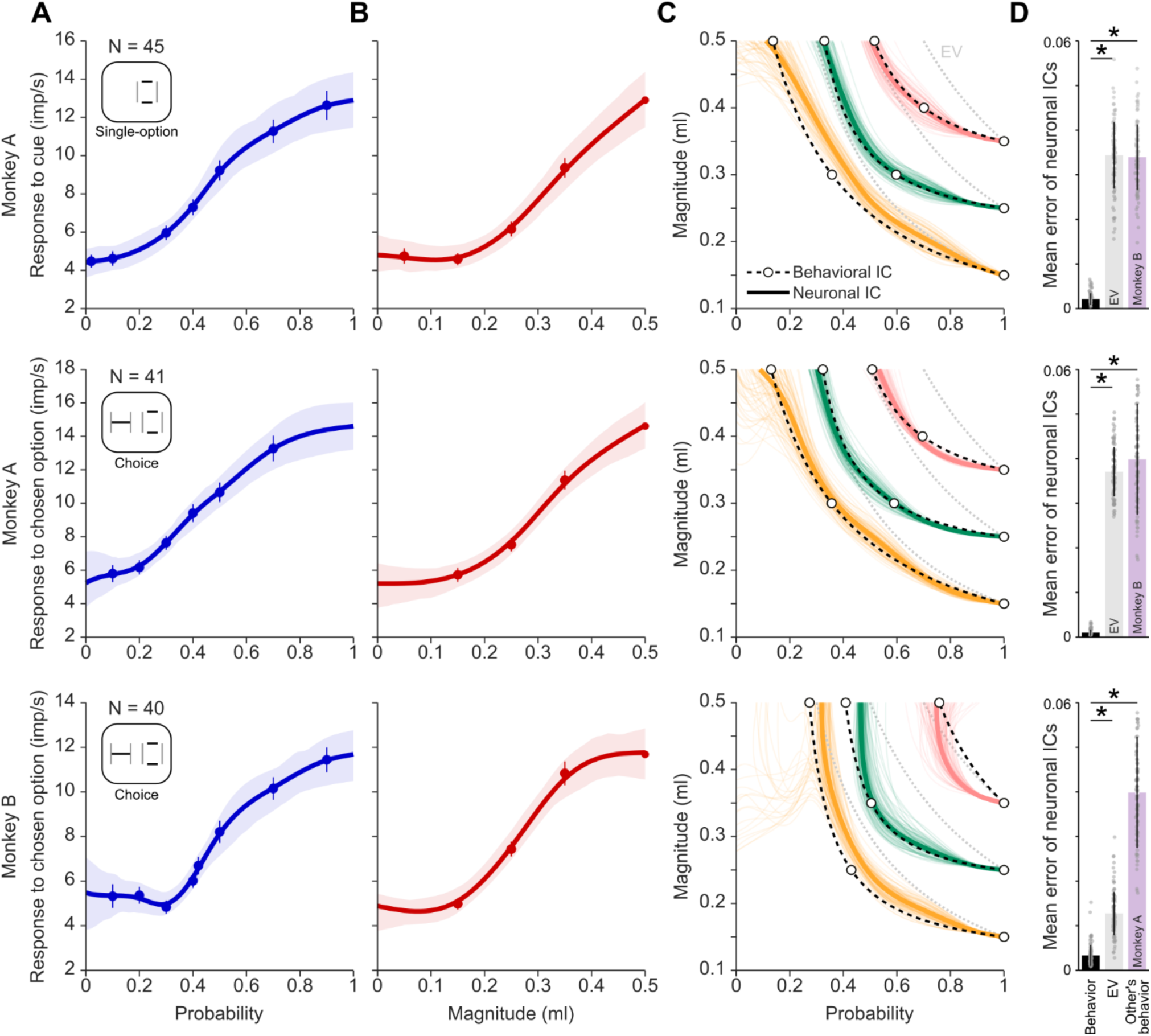
Neuronal indifference curves match behavioral indifference curves: evidence for subjective value derived from nonlinear probability and magnitude responses. A) Nonlinear regression of population responses to gamble probability. Average response (dots), SEM (bars) and 95% confidence interval of spline regressions (area) were computed across the bootstrapped distribution (1,000 resamples with replacement) of population responses. Gamble magnitude was fixed at 0.5 ml. B) Nonlinear regression of population responses to safe option magnitude. Conventions as in panel A. C) Neuronal indifference curves (ICs). The nonlinear magnitude and probability responses were combined multiplicatively to obtain a neuronal value measure for all gambles in the tested range. ICs were defined as points with the same neuronal value measure. The neuronal ICs were computed represented an out-of-sample value measure as points within the D) Deviation of neuronal ICs from behavior. Mean distance (across all IPs) between neuronal ICs and behavioral ICs (black), equal-EV curves (gray), and other animal’s behavioral ICs (purple). Average (bars) and standard deviation (vertical lines) were computed from the bootstrapped resamples (dots, N = 100). The neuronal ICs were closer to the behavioral ICs than to the equal-EV curves or to the other animal’s ICs (Kolmogorov-Smirnov test, asterisk: P < 1E-32, N = 100). See Figure S4 for detailed analysis.

We combined these nonlinear functions in a multiplicative manner, to compute a putative neuronal economic subjective value measure for every gamble in the tested range (resolution 0.01, see Methods). Importantly, this was computed separately for each animal to retain differences in evaluation between subjects and considering that utilities cannot be theoretically compared between individuals. We then used this value map to construct neuronal indifference curves (ICs), defined as points with equal subjective value. Equivalent to the behaviorally defined ICs, we constructed three ICs starting from safe options with magnitudes 0.15, 0.25 and 0.35 ml respectively. The resulting neuronal indifference map captured in high detail the shape of the behavioral indifference map in both single-option and choice tasks (Figure 5C). The neuronal ICs closely matched the animal-specific behavioral ICs, differing from objective equal-EV curves (Figure 5D). Note that the ICs computed from the activity of one animal’s dopamine neurons did not match the other animal’s ICs (Figure S4). These results support the hypothesis that neuronal subjective values stem from the integration of subjectively weighted reward magnitude and probability components.

To evaluate the effective contribution of nonlinear probability coding to value computations, we repeated the same analysis assuming linear probability coding (Figure 6). In this simplified model, subjective values coincided with expected utilities, as defined in EUT. Linear probability weighting produced ICs that did not overlap with the behavioral curves (Figure 6C), resulting in a significantly larger neuronal-behavioral difference compared with the nonlinear probability-weighting model (Figure 6D) (Kolmogorov-Smirnov test on the distribution of bootstrapped mean squared errors, P < 1E22, N = 100). This result indicated that the nonlinear coding of probabilities significantly contributed to the encoding of subjective values. Neuronal ICs computed assuming an additive computation (i.e., value calculated as the sum of magnitude and probability information) did not match behavioral ICs as well as the multiplicative model (Figure 6D, Figure S6).

**Figure 6.**
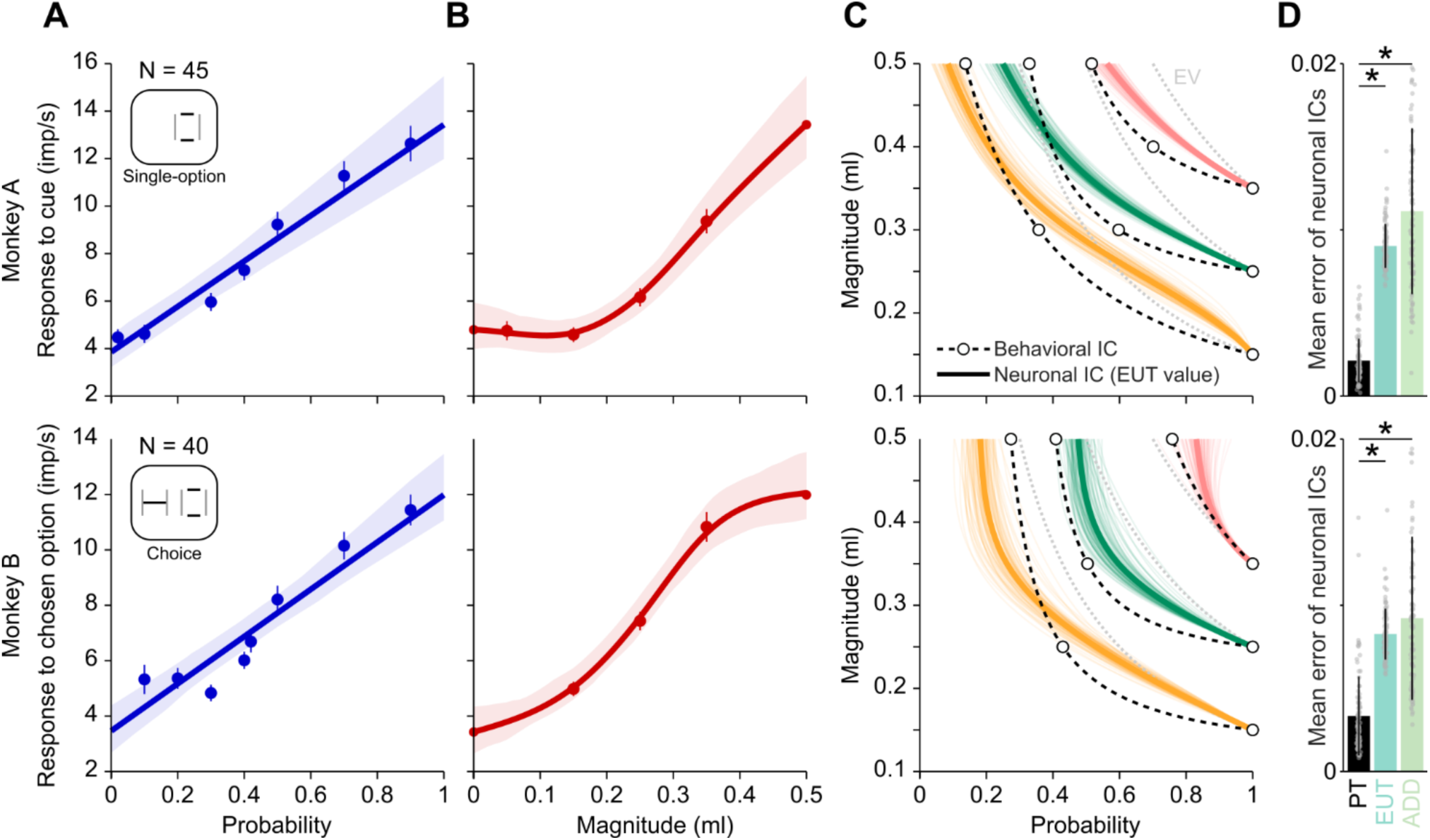
Linear probability assumption prevents subjective value coding, as evidenced by reduced neuronal-behavioral match. A) Linear regression of neuronal population responses to gamble probability. Conventions as in Figure 5A. B) Nonlinear regression of population responses to safe magnitude. Conventions as in Figure 5B. C) Neuronal indifference curves (ICs) computed assuming linear probability coding did not match the behavioral ICs. Conventions as in Figure 5C. D) Deviation of neuronal ICs from behavior for different assumptions for value computation. The mean error (mean squared distance between neuronal and behavioral IPs along the probability dimension) was computed assuming either nonlinear probability weighting as in prospect theory (PT), linear probability weighting as in expected utility theory (EUT) or the additive combination of neuronal utility and weighted probability (ADD). The PT value computation resulted in smaller neuronal-behavioral difference compared to both EUT and ADD assumptions (Kolmogorov-Smirnov test, P < 1E22, N = 100). Equivalent results were obtained in the single-option task (top, monkey A) and during choice (bottom, monkey B; see Figure S5 for monkey A’s choice data).

Overall, these data show that dopamine neurons’ signals reflect the nonlinear weighting of reward magnitude and probability, the two fundamental economic constructs forming the basis for subjective value computation. Dopamine signals thus reflect the integration of both subjective reward magnitude (utility) and subjectively weighted reward probability into economic value, as shown by the faithful neuronal reconstruction of the animal-specific indifference maps elicited from choice.

### Dopamine responses reflect choice biases violating the independence axiom of EUT

To finely quantify the effect of nonlinear probability weighting on behavior, we defined a test based on the independence axiom of EUT, which is violated if probabilities are not linearly evaluated. Systematic independence axiom violations have been reported in humans (Allais, 1953; Kahneman and Tversky, 1979; Starmer, 2000) and monkeys (Ferrari-Toniolo et al., 2022). We quantified axiom violations from both behavioral and neuronal data and assessed the correlation between the two quantities. As an out-of-sample neuronal test, we compared the behavioral violations with those predicted from neuronal responses in different sets of trials.

According to the independence axiom of EUT, preferences between two options should not change after adding the same probability of a common outcome to both options. For testing the axiom’s predictions, we defined two option pairs ({A, B} and {C, D}), with options C and D constructed via a common transformation of options A and B, respectively (Figure 7A). Preferring A and C (or B and D) complies with the axiom, while preferring A and D (or B and C) violates it. In the Machina triangle, a scheme in which points are gambles with probabilities associated to three possible outcomes (0, 0.25 and 0.5 ml), the {A, B} and {C, D} option pairs appear as parallel lines (Figure 7B). EUT implies linear and parallel indifference curves within the Machina triangle, thus predicting matching preferences between the two option pairs. On the other hand, nonlinear probability weighting produces nonlinear indifference curves, predicting different preferences between the two option pairs (i.e., a preference change), which violates the independence axiom.

**Figure 7.**
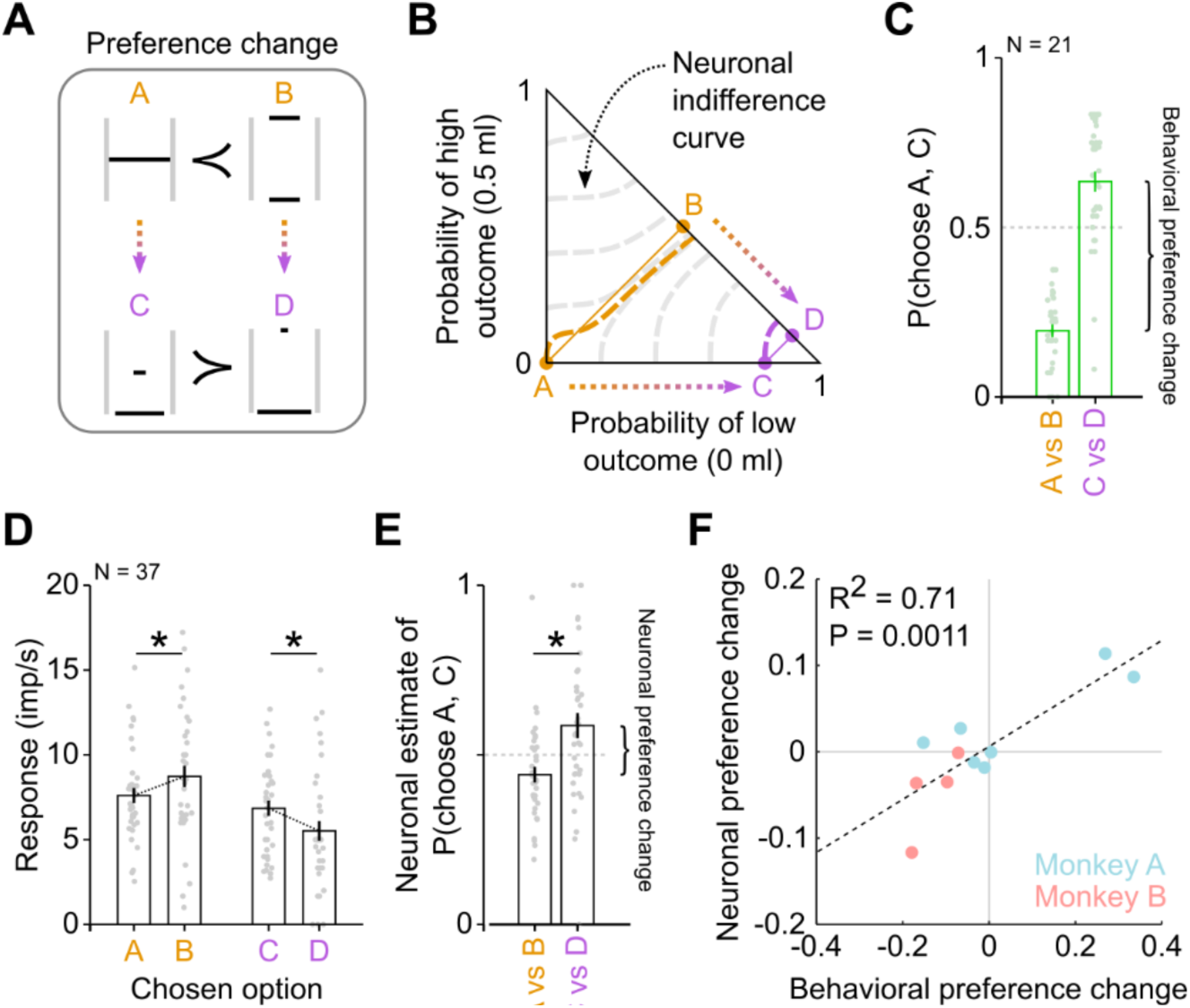
Dopamine responses reflect choice biases predicted with nonlinear probability weighting. A) Choice options used for measuring preference change. Options C and D are constructed from options A and B respectively, by scaling the probability of the high outcome by the same factor. According to EUT, preferences should not change between the {A, B} and {C, D} option pairs. Nonlinear probability weighting can result in preference changes (common ratio effect). B) Neuronal indifference curves predict preference change. Scheme of independence axiom test in the reward probability space (Machina triangle), in which each point correspond to a gamble with a specific combination of probabilities associated to three possible outcomes (0, 0.25 and 0.5 ml). According to EUT, indifference curves (points with equal subjective value) within the Machina triangle are linear and parallel, while being curved when probabilities are nonlinearly weighted (as implemented in cumulative prospect theory). Indifference curves estimated from the dopamine magnitude and probability responses (see Methods for details) were curved and “fanning-in”, predicting a change in preferences between {A, B} and {C, D}. Data from Monkey B. C) Behavioral preferences. Monkey B preferred option A to B, while preferring option C to D. This choice pattern cannot be explained by the linear-probability model, violating the independence axiom of EUT. Dots: single-session preference (probability of choosing option A, C). Bar height: average preference across recording sessions (N = 21). Vertical lines: SEM. D) Responses of dopamine neurons to the four options reflect behavioral preferences. Responses to option B were significantly higher than to option A (P = 0.024), while responses to option C were significantly higher than to option D. Dots: average response of one neuron across all trials in which the corresponding option (horizontal axis) was chosen. Bar height: average response across neurons (N = 37). Vertical lines: SEM. E) Neuronal estimated preferences. Neuronal preference for one option were estimated as the percentage of responses being higher for that option than for the alternative option. F) Significant correlation between behavioral and neuronal preference change. Across different tests (using different sets of options; see Figure S7), neuronal-estimated preference changes reflected behavioral preference changes.

For a subset of dopamine neurons (N = 32 in Monkey A; N = 37 in monkey B), we offered the monkey choices between the axiom-based {A, B} and {C, D} options, in addition (and intermingled with) choices between different reward magnitude and probability levels. For an out-of-sample neuronal prediction of axiom violations, we estimated indifference curves within the Machina triangle from the neuronal responses and predicted the corresponding choice pattern. These neuronal indifference curves, computed from the dopamine responses to different magnitude and probability levels (see Methods), appeared markedly curved (Figure 7B, dashed). Their “fanning-in” pattern predicted preferences violating the independence axiom, with options B and C lying on higher indifference curves (i.e., higher subjective values) compared to options A and D, respectively. Behavioral choices confirmed this prediction, with option B and C chosen over options A and D in the majority of trials (Figure 7C).

As a direct test for subjective value coding during choices violating the axiom, we examined the dopamine responses to the four options. The average population response was higher for option B than for option A (t test, P = 0.024), while also being higher for option C than for option D (P = 0.014) (Figure 7D), reflecting behavioral preferences. A neuronal estimate of the choice probability (as proportion of responses higher for one option than for the alternative), confirmed a behavior-matching preference change between the two option pairs (Figure 7E). Across different sets of options (N = 11 for monkey A; N = 4 for monkey B; see examples in Figure S7), we found a significant correlation between neuronal and behavioral preference changes (Figure 7F). Note that monkey A neurons reflected the animal’s mostly positive preference change, while monkey B neurons reflected the animal’s negative preference change.

These results show that, when choices are biased in a specific direction, as in violations of EUT’s independence axiom, dopamine responses closely reflect the behavioral choice biases. The correct prediction of axiom violations solely from the nonlinear dopamine responses highlights the accurate neuronal value code resulting from the subjective integration of magnitude and probability information.

### Heterogeneous coding of reward components across dopamine neurons

Our previous data analyses indicated the meaningful coding of subjective reward components in a population of dopamine neurons. The averaged population activity leaves an open question about the contribution of individual neurons to the subjective value computation. Furthermore, recent studies have identified heterogeneity across dopamine neurons’ value responses (Dabney et al., 2019). We next investigated the degree of heterogeneity in the coding of value components across neurons.

We characterized the responses of individual neurons to different reward magnitude and probability levels. To this aim, we curve-fitted the response profiles using spline functions, identifying different response patterns in the magnitude and probability domains. We found dopamine neurons showing convex or concave responses to both magnitude and probability, while other neurons showed a mixture of convex and concave nonlinearities across the two dimensions (three example neurons in Figure 8A). As the shapes of utility and weighted probability define subjective values, the different convexity in individual neurons’ responses potentially represented different subjective values and, correspondingly, different choice patterns and risk attitudes. To quantitatively characterize the neurons’ response convexity, we calculated the relative scaling factor (RSF) as defined in a previous study (Dabney et al., 2019). In brief, we subtracted the pre-cue activity from the responses, which resulted in a transformed response pattern switching from negative to positive values (reversal point). We then performed two linear regressions (before and after the reversal point) and computed the RSF as the slope-ratio between the two regression lines. The RSF indicated, in a simple scalar metric, convex (RSF > 0.5) or concave (RSF < 0.5) shapes. The resulting RSF varied broadly across the neuronal population (Figure 8B), confirming previous observations. Interestingly, we found a significant correlation between the RSFs computed separately for magnitudes and for probabilities (Pearson’s R^2^ = 0.20, P = 3.3e-4) (Figure 8C), indicating that individual dopamine neurons had generally similar response shapes across the magnitude and probability domains. The reversal point in the magnitude domain also significantly correlated with the reversal point in the probability domain (Pearson’s R^2^ = 0.16, P = 1.3e-3) (Figure 8D).

**Figure 8.**
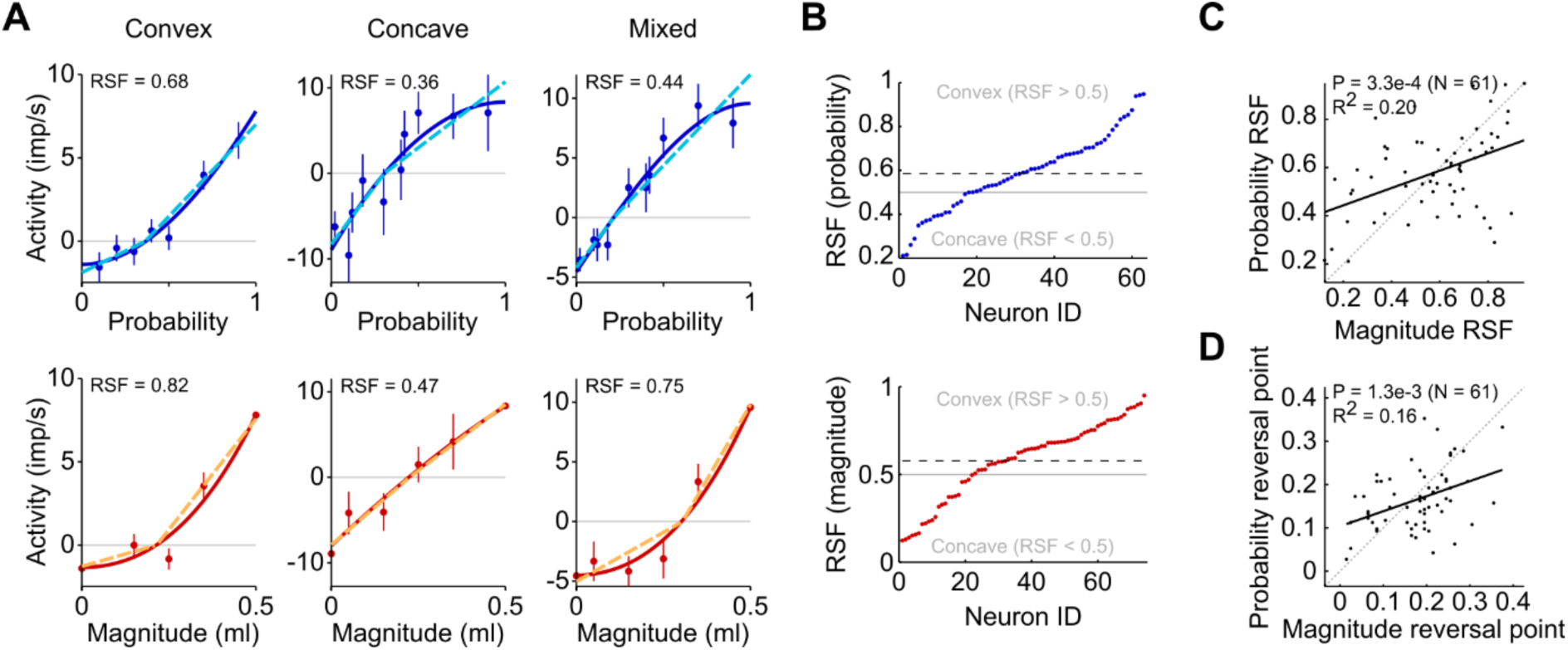
Heterogeneous subjective value coding across individual dopamine neurons. A) Three example neurons highlighting different nonlinear responses to reward probabilities (top) and magnitudes (bottom). Baseline-subtracted activities (dots: mean across trials; bars: SEM) were modelled with spline functions (continuous curves) as well as with two linear regressions (dashed lines) separately for negative and positive values (as in Dabney et al., 2019). The two methods similarly captured the variability in nonlinear value coding across neurons. The relative scaling factor (RSF), i.e., the slope-ratio between the two regression lines, indicated convex (RSF > 0.5) or concave (RSF < 0.5) shapes. B) The broad spectrum of RSF values across neurons (data from both monkeys) highlighted the value-coding heterogeneity for responses to both probability (top) and magnitude (bottom), suggesting heterogeneous weighted probability and utility profiles in single neurons. Dashed line: mean RSF across neurons. Neurons sorted by increasing RSF values, independently for probability and magnitude. C) Correlation between probability-based and magnitude-based RSF. D) Correlation between probability-based and magnitude-based reversal points (i.e., probability/magnitude values at which the activity crossed zero).

These data indicate a degree of variability in the coding of utility and subjectively weighted probability across individual dopamine neurons. Despite this heterogeneity, the average activity across these heterogeneous neurons resulted in a value code accurately describing the monkey’s choice behavior and risk attitudes.

## DISCUSSION

Our data show that primate dopamine neurons encode subjective economic values resulting from the integration of subjectively transformed reward magnitudes and probabilities. We operationally defined subjective values from the continuity axiom of EUT, resulting in a behavioral indifference map which captured the subjective pattern of choices and risk attitudes (Figure 1). Guided by the same principle, we constructed a neuronal indifference map, demonstrating a fine match between the neuronal and behavioral measurements (Figure 2 and 3). Crucially, the dopamine value signal reflected individual preferences regardless of the physical composition of the choice options (Figure 4). Analyzing responses to magnitude and probability separately, we isolated nonlinearities in the dopamine value code which reflected the subjective transformations of the two reward components (Figure 5). In particular, the contribution of nonlinear probabilities appeared essential in reconstructing animal-specific, behavior-compatible economic values (Figure 6 and 7). Thus, despite heterogeneous value coding in individual neurons (Figure 8), the dopamine population signal conformed with the integration of subjectively weighted magnitude (utility) and subjectively weighted probability, reflecting the animal’s choices, including their specific risk attitudes and choice biases. These data demonstrate an implementation of the theoretical construct of subjective value in the brain that has been used for explaining economic choices for more than 50 years now.

Subjectively weighted probability is a critical components for the evaluation of choice options (Allais, 1953; Kahneman and Tversky, 1979; Gonzalez and Wu, 1999), along with utility, temporal discounting and satiety (von Neumann and Morgenstern, 1944; Frederick et al., 2002; Rolls et al., 1983). Weighted probability was introduced to explain unexpected choice patterns in humans, including violations of EUT’s independence axiom (Starmer, 2000). It crucially contributes to value computations, resulting in the expression of individual risk attitudes. Dopamine signals are compatible with the coding of utility (Stauffer, Lak and Schultz, 2014) and temporal discounting (Kobayashi and Schultz, 2008). By now identifying subjectively weighted probability coding in dopamine neurons, we provide the crucially missing component of a plausible brain mechanism for computing subjective values.

Our results show that dopamine neurons represent the integration of different reward components. Open questions remain about the neural circuits and mechanisms that contribute to this integrated value signal. One possibility is that the objective reward components are separately transformed into subjective quantities which then converge into the integrated dopamine value signal. This hypothesis implies the existence of different sets of neurons, possibly in different brain regions, encoding the subjective representations of each reward component separately. Another fundamental question regards the formation and development of the reward magnitude and probability nonlinearities. The specific shapes of the economic functions may result from the efficient allocation of limited neural resources (Polanía et al., 2019; Woodford, 2012; Zhang et al., 2020). They may also be influenced by past rewards, as evidenced by our previous observation of economic functions adapting to the reward history (Ferrari-Toniolo et al., 2019; Bujold et al., 2021). It remains to be investigated whether dopamine value signals reflect these adaptations.

When two choice options were presented simultaneously, dopamine neurons encoded the subjective value of the option that was selected later in the trial. This type of response qualifies as chosen value signal, which could reflect either the option’s value averaged over several trials, or the trial-by-trial value fluctuations driving individual-trial preferences. A recent study supported the latter mechanism, with dopamine neurons encoding the trial-by-trial values that were explicitly indicated by monkeys during an auction task, despite constant reward amounts (Hill et al., 2024).

We report a degree of heterogeneity in the representation of subjective value components across individual dopamine neurons in primates, which extends previous findings in rodents concerning heterogeneity in the slopes of reward prediction error responses (Dabney et al., 2019). Furthermore, we found significant correlation between the variability of the magnitude and probability components, suggesting that the heterogeneity predominantly affected the overall combined subjective value rather than the individual reward components. Our pooling of heterogeneous responses was justified by their match with behavior, which was obtained without any arbitrary selection of neuronal populations. Nevertheless, the observed heterogeneity could be suitable for projecting different value signals to different brain structures (Morales and Margolis, 2017; Beier et al., 2015; Lammel et al., 2008). In line with this hypothesis, a recent study showed that the stimulation of different dopamine pathways produced different behavioral risk attitudes (Sasaki et al., 2024).

Other brain regions have been associated with economic value coding. Value signals in the primate orbitofrontal cortex (OFC) are causally related to choice (Ballesta et al., 2020). We showed in a previous study that OFC signals integrate reward magnitude and probability (Ferrari-Toniolo and Schultz, 2023), although only an OFC subpopulation encoded behavior-matching economic subjective value. By comparison, OFC neurons varied more in their value code than dopamine neurons. Different components of prospect theory (PT), including loss aversion and reference points, were used to characterize neuronal responses in the insular cortex (Yang et al., 2022), striatum and OFC (Imaizumi et al., 2022), suggesting the distributed coding of PT-compatible reward variables across brain regions. In addition to value, neurons encoding risk (i.e., variance in the reward distribution) were observed in the amygdala (Grabenhorst and Baez-Mendoza, 2024) as well as in OFC (O’Neill and Schultz, 2010). These more abstract reward-related neuronal representations, which may contribute to the formation of subjective values and guide choices in particular situations, were not observed in our current study: dopamine neurons appeared to encode the overall subjective economic value, integrated across the underlying reward components of utility and weighted probability.

## ACKNOWLEDGEMENTS

The work was supported by a Wellcome Trust Principal Research Fellowship to WS, by Wellcome Grants WT 095495 and WT 204811, and by European Research Council (ERC) Advanced Grant 293549. We thank Aled David and Christina Thompson for animal and technical support, Dr. Polly Taylor for expert anesthesia, Dr. Henri Bertrand for veterinary support. For the purpose of Open Access, the authors have applied a CC BY public copyright license to any Author Accepted Manuscript version arising from this submission.

## AUTHOR CONTRIBUTIONS

Experimental design, SF-T and WS; Investigation, SF-T, LCUS; Data Analysis, SF-T; Writing and Editing, SF-T and WS.

## SUPPLEMENTARY INFORMATION

**Figure S1.**
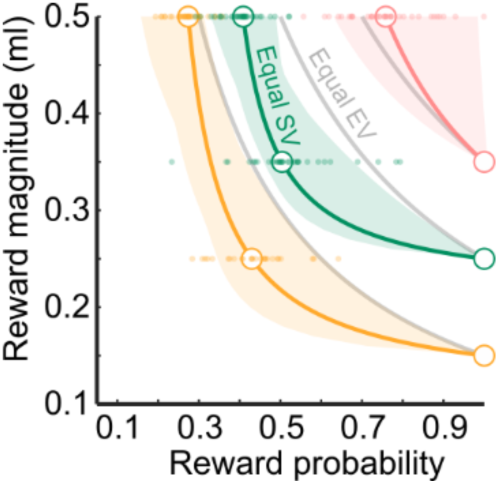
Behavioral indifference map. Indifference curves estimated from 35 behavioral sessions in monkey B. See Figure 1D for full details.

**Figure S2.**
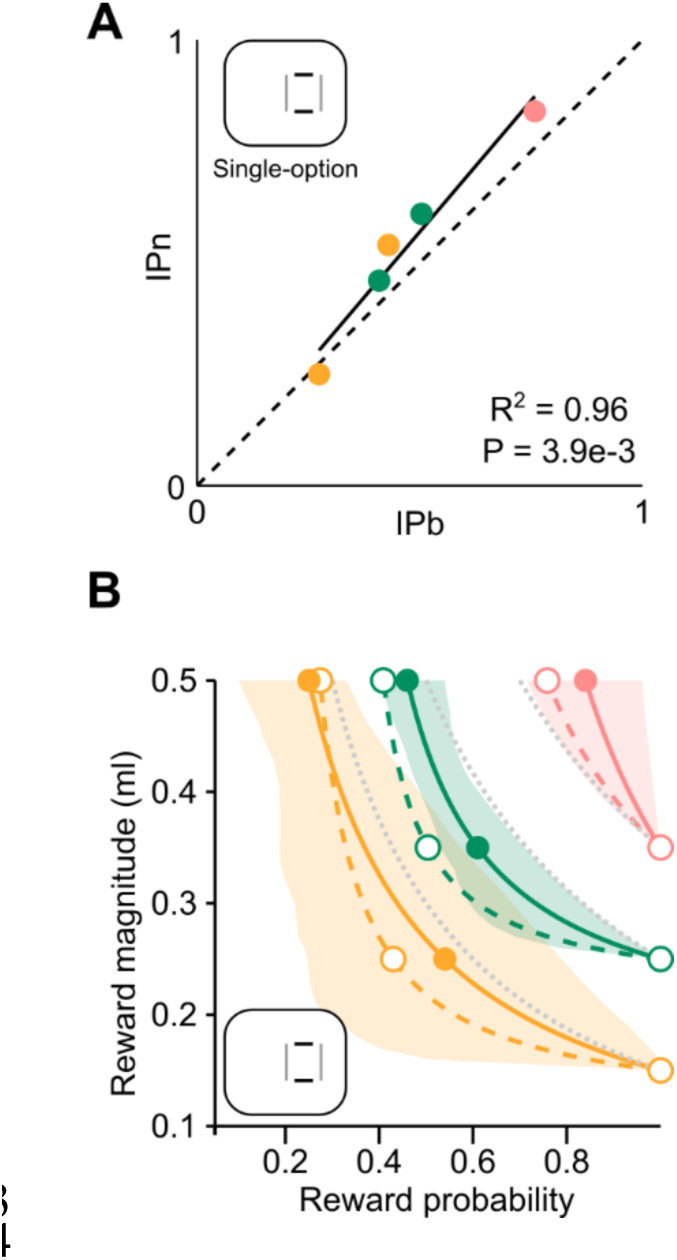
Neuronal-behavioral match of indifference points. Data from monkey B. Correlation between neuronal and behavioral subjective value measures (panel A) and comparison of neuronal and behavioral indifference curves (panel B). See Figure 3 for full details.

**Figure S3.**
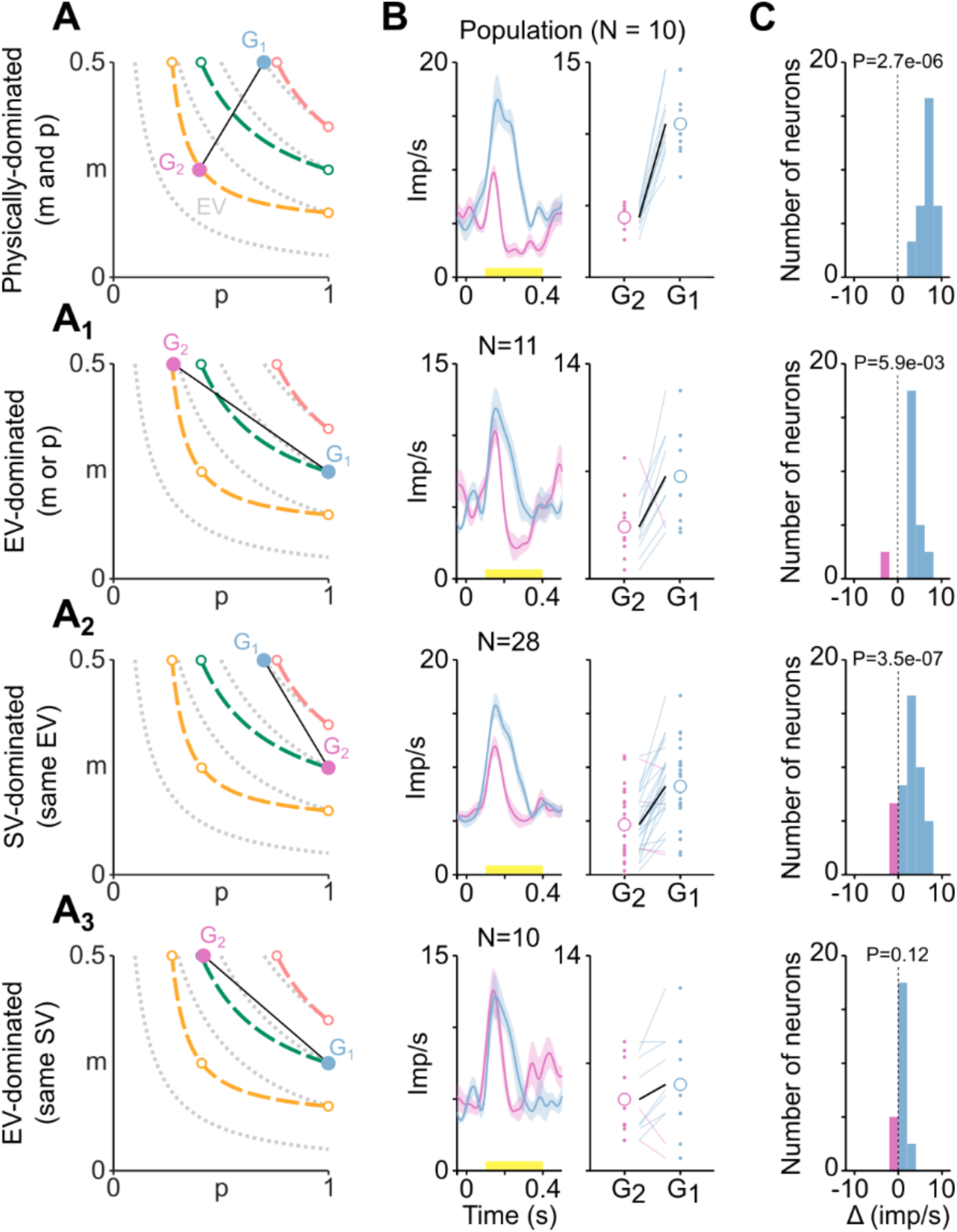
Neuronal responses reflects subjective value rather than physical value. Data from monkey B, single-option task. See Figure 4 for full details.

**Figure S4.**
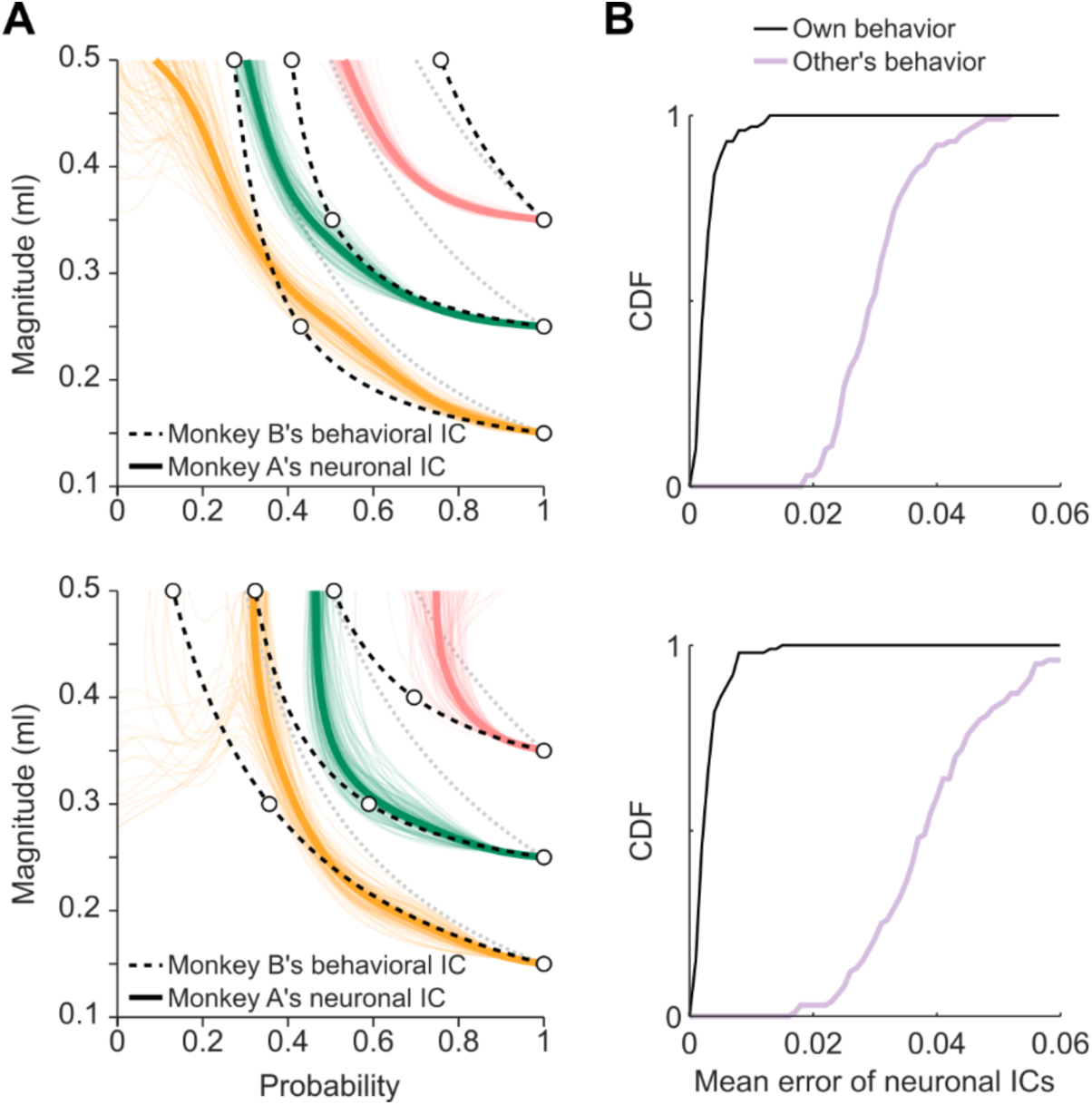
Responses of one monkey’s dopamine neurons did not match the other monkey’s behavior. A) Neuronal indifference curves (ICs) differed from the behavioral ICs of the other monkey. Neuronal ICs computed from monkey A’s dopamine responses (top, monkey B in bottom panel), compared with behavioral ICs computed from monkey B’s behavior (top, monkey A in bottom panel). See Figure 5C for full details. B) Distance between neuronal and behavioral ICs for own and other’s behavior. Cumulative distribution function (CDF) of the mean neuronal-behavioral ICs errors was computed in the probability (horizontal) dimension and averaged across all IPs commonly measured in the two animals. The error measure was repeated for 100 resamples (with replacement), obtaining the represented distribution of errors. The mean error distribution for other’s behavior was significantly different from that of the own behavior (Kolmogorov– Smirnov test, P < 1E-44, N = 100).

**Figure S5.**
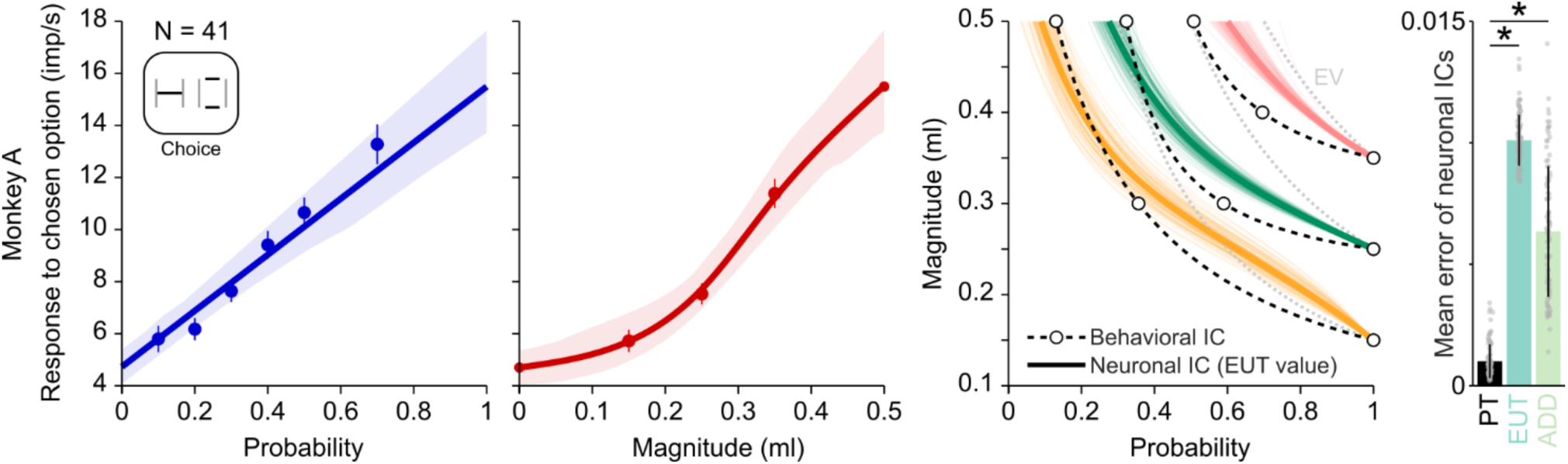
Linear probability assumption prevents subjective value coding. Data from monkey A in the choice task. See Figure 6 for full details.

**Figure S6.**
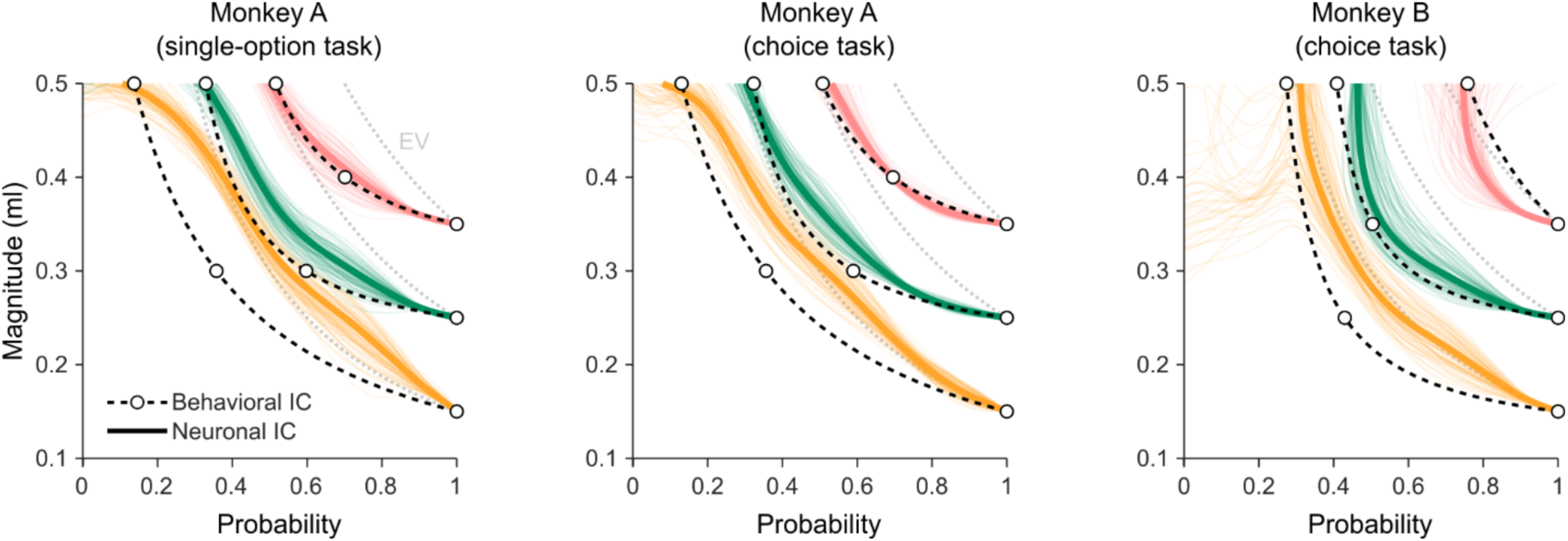
Additive combination of neuronal magnitude and probability responses. Neuronal ICs computed assuming additive value coding did not match the behavioral ICs as well as with the multiplicative value assumption. The discrepancy is especially evident for lower ICs. Conventions and symbols as in Figure 6. See Figure 6 and Figure S5 for statistical comparisons between additive (ADD) and multiplicative value computations.

**Figure S7.**
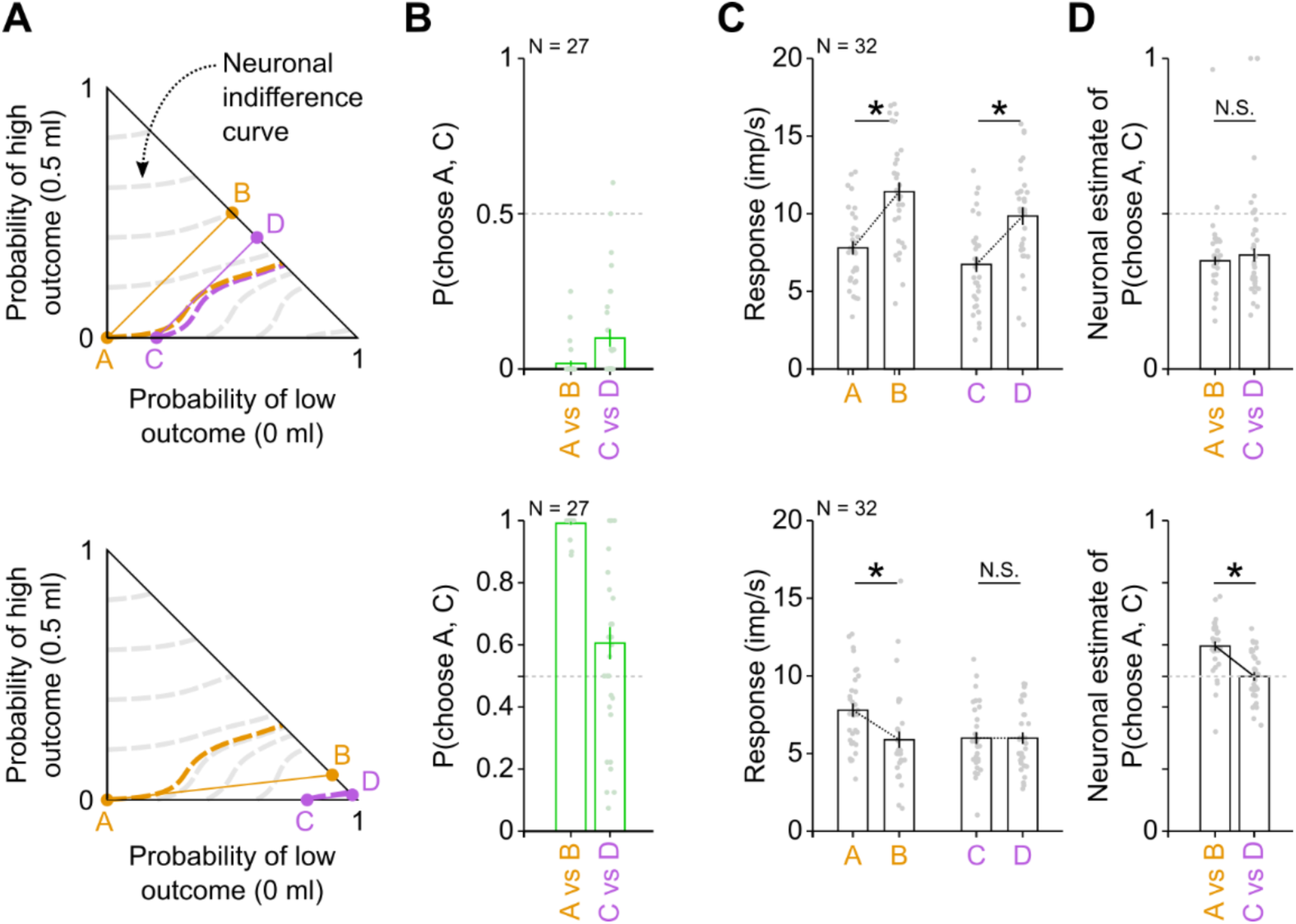
Independence axiom compliance and violation, reflected in dopamine responses. Examples of behavioral compliance (top) and violation (bottom) of the independence axiom of EUT, reflected in dopamine responses. Neuronal indifference curves (panel A), which were computed out-of-sample from magnitude and probability responses, predict axiom compliance for specific option sets (top) and axiom violation for other option sets (bottom). This violation pattern was confirmed in behavioral tests (panel B), which showed strong preference for B and D (no violation) in the first options set, and a change in preference in the second options set (A strongly preferred to B, while C similarly preferred to D). The population of dopamine neurons (N = 32) reflected measured preferences (panel C) and options-set-specific preference changes (panel D). Data from monkey A in choice task. Full details in Figure 7B-E.

## ONLINE METHODS

### Experimental models and subject details (Rhesus monkey, *Macaca mulatta*)

Two adult male macaque monkeys (monkey A, “Aragorn”; monkey B, “Tigger”), weighing 11 kg and 13 kg, respectively, were used in the experiments. The animals were born in captivity at the Medical Research Council’s Centre for Macaques (CFM) in the UK. Both animals had not been used in other studies.

The animals had been surgically implanted with a headpost and a recording chamber for neurophysiological recording. They were headposted for 2 – 3 h on each test day of the current experiment. Both animals had previous experience with the visual stimuli and experimental setup (Ferrari-Toniolo et al., 2019, 2022).

All experimental procedures had been ethically reviewed and approved and were regulated and continuously supervised by the following institutions and individuals in the UK and at the University of Cambridge (UCam): the Minister of State at the UK Home Office, the Animals in Science Regulation Unit (ASRU) of the UK Home Office implementing the Animals (Scientific Procedures) Act 1986 with Amendment Regulations 2012, the UK Animals in Science Committee (ASC), the local UK Home Office Inspector, the UK National Centre for Replacement, Refinement and Reduction of Animal Experiments (NC3Rs), the UCam Animal Welfare and Ethical Review Body (AWERB), the UCam Governance and Strategy Committee, the Home Office Establishment License Holder of the UCam Biomedical Service (UBS), the UBS Director for Governance and Welfare, the UBS Named Information and Compliance Support Officer, the UBS Named Veterinary.

### Experimental design

Each animal made choices between a gamble and a safe option. Options were mutually exclusive and collectively exhaustive and appeared simultaneously and at equal distance from the animal in front of it on a computer monitor (eye-monitor distance: 50 cm). All gambles had one non-zero outcome, which could vary in magnitude (m, from 0 ml to 0.5 ml in 0.05 ml steps) and probability (p, from 0 to 1; resolution: 0.02) and were represented respectively by the vertical position and width of a horizonal bar (Figure 1A, B). A safe option represented a degenerate gamble (p = 1). For each option, we set the magnitude and probability of reward independently. The bar’s horizontal position was randomly shifted on each trial (within the two vertical bars) to avoid delivering information about the expected value (EV) of the gamble (a fixed bar’s edge position would completely identify the gamble’s EV). The learning of the stimuli involved an initial training of the quantitative stimuli, varying one of the variables over two or three levels, and did not include the full stimulus variety which the animals generalized from their initial training experience.

A second type of options was introduced later in the training, with gambles including three possible outcomes. The new visual cues were consistent with the previously learned cues, representing three-outcome gambles with three horizontal bars (vertical position: outcome magnitudes; horizontal width: outcome probability). In this task, option pairs included one- or two-outcome gambles versus two- or three-outcome gambles. The magnitudes of the three outcomes were fixed (0, 0.25 and 0.50 ml) while their probabilities varied (from 0 to 1; resolution: 0.02).

A trial started when a cross appeared at the center of the computer monitor (Figure 1A). A cursor, controlled by the monkey through left-right joystick hand movements, was required to be screen-centered, and after a control period (1 – 1.5 sec) two visual Cues appeared, representing the two choice options, whose fixed left and right positions alternated pseudorandomly. This alternation ensured that any potential side bias would not affect our analyses on the average choices. We inferred subjective reward value from observable choice between two options. Thus, the animal revealed its preference at the time of choice. Such “revealed preferences” contrast with “stated preferences” that are expressed, typically in humans, at some interval or distance from the actual choice and therefore reflect subjective value in a less reliable manner. In our experiment, the monkey revealed its preference by moving the cursor toward the chosen option. After holding it there for 1 sec, a liquid reward (water) was delivered through a spout directly in front of the animal’s mouth. The reward was contingent on the selected option’s magnitude and probability. Each option pair was presented for 6 to 12 replications. All option pairs were pseudorandomly interleaved.

We estimated subjective economic values at choice indifference between a gamble with varying reward probability and a fixed safe reward. Tests at choice indifference are immune to slope differences of choice functions that might confound value estimation. An option was inferred to have the same subjective value as its alternative when both options were chosen with equal probability (P = 0.5 each of two rewards in binary choice). To find the gamble producing equal choice probability as the fixed safe option, we held the safe reward magnitude constant (reward probability p = 1.0) and varied the reward probability of the gamble option across the full probability range of 0 and 1. We estimated choice indifference from an S-shaped psychophysical choice function (softmax, see Eq. 2 below) fitted to the probabilities of choosing the gamble over the safe option (Figure 1C). We plotted two-dimensional indifference points (IP) by positioning each equally frequently chosen option with its specific reward magnitude and probability at the intersection of the x-coordinate (probability) and y-coordinate (magnitude) of a two-dimensional plot. Irrespective of their different magnitude-probability composition, the IPs of all equally frequently chosen options align as an indifference curve (IC) (colored curves in Figure 1D). As we inferred subjective economic value from observable choice, we assumed that all options on the same IC have the same subjective economic value, and options on different ICs have different value. Options on higher ICs (farther from origin) are more frequently chosen than options on lower ICs, and thus have higher value. For example, options on the pink IC in Figure 1D is more frequently chosen, and have higher value, compared to options on the amber and green ICs.

All data collection and analyses were performed using custom code in MATLAB (version 8.3.0 (R2014a). Natick, Massachusetts: The MathWorks Inc).

### Using the continuity axiom for value estimation

Expected Utility Theory (EUT) proposes four axioms that determine the maximization of utility (von Neumann and Morgenstern, 1944). Compliance with completeness (axiom I) and transitivity (axiom II) is necessary for consistently ranking all choice options. Compliance with continuity (axiom III) demonstrates that choices reflect a meaningful representation of numerical utility: the continuity axiom implies the existence of a value function. The independence axiom (axiom IV) defines the computation of expected utility from reward magnitudes and their probabilities. The four axioms of EUT define necessary and sufficient conditions for choices to be described by the maximization of subjective economic value: if the axioms are fulfilled, a subjective economic value can be computed from the reward components (magnitude and probability) for each choice option, and the decision maker behaves as if choosing the highest valued option. Violations of the independence axiom, reported in both human and animal subjects (Allais, 1953; Kagel et al., 1990; Ferrari-Toniolo et al., 2022), led to the development of non-expected utility theories, such as prospect theory (PT) (Kahneman and Tversky, 1979), which required a weakened form of the independence axiom while retaining the continuity axiom requirement.

The continuity axiom can be formally stated as:

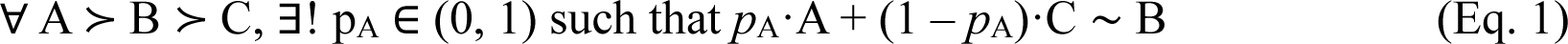

where A, B, C and AC are gambles, “≻” is the preference relation, and “∼” the indifference relation. In words, compliance with the continuity axiom requires to identify a reward probability *p*_A_ at which choice indifference occurs between a fixed gamble B and a variable gamble AC. The variable gamble AC consists of a probabilistic combination of a higher valued gamble A (with probability *p*_A_) and a lower valued gamble C (with probability 1 - *p*_A_) (Eq. 1). In our experimental design, we used sure rewards (p = 1) as options A, B and C, resulting in binary choices between a fixed safe option (B) and a gamble with variable probability and fixed magnitudes (corresponding to the AC option in Eq. 1). We varied reward magnitudes of A and B between 0.05 ml and 0.5 ml, while C was kept fixed at 0 ml (i.e., the gamble had one non-zero outcome). The continuity axiom with this extended testing scheme constitutes the foundation of the current study.

The behavioral indifference point (IPb) represents a utility measure, being a numerical quantity associated with the subjective evaluation of reward B in relation to outcomes A and C. As a deterministic rule, the axiom assumes constant preferences over time. We interpreted the axiom in a stochastic sense, measuring preferences stochastically over repeated choices, which can potentially fluctuate over time. The IPb was computed according to a standard discrete choice model: we fitted a softmax function to the probability of choosing the AC option and identified the point for which the softmax’ value was 0.5, which corresponded to a 0.5 probability of choosing equally frequently each option, i.e., choice indifference (Figure 1C). The following softmax function was fitted to the choice data through non-linear least squares (Matlab function: *nlinfit*):

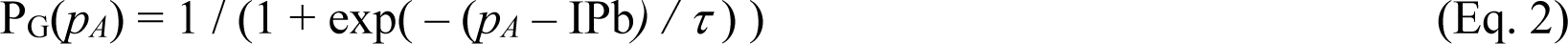

With P_G_ representing the proportion of gamble choices, IPb corresponding to the probability *p_A_* resulting in equal preference for the two options, and *τ* as softmax “temperature” parameter, representing the steepness of the choice function (steeper for lower τ values), in analogy to the Boltzmann distribution in statistical mechanics. In alternative formulations, the softmax steepness is referred to as β (“inverse temperature” or “precision”), i.e., the reciprocal of the temperature parameter defined here.

Behavioral indifference curves (i.e., series of IPb’s computed for the same fixed safe option) were computed by fitting a parametric function to the IPb’s (Matlab function: *nlinfit*). We used the following hyperbola equation as parametric function:

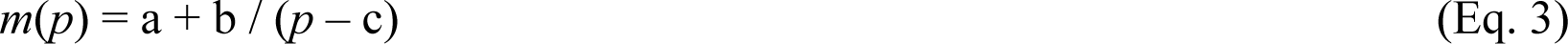

with *m* representing the reward magnitude, *p* the reward probability, and a, b, and c as free parameters (initial values set to 0, 0.1, 0 respectively).

### Neurophysiological recordings

A head-restraining device and a recording chamber were implanted on the skull under full general anesthesia and aseptic conditions. The chambers were 35 mm wide by 40 mm long (Gray Matter). The stereotactic coordinates of the chamber enabled neuronal recordings from the dopaminergic midbrain and orbitofrontal cortex. We located the midbrain dopamine region (right hemisphere) from bone marks on coronal and sagittal radiographs taken with a guide cannula inserted at a known coordinate in reference to the implanted chamber.

We conducted single-neuron electrophysiological recordings using glass-coated tungsten electrodes, custom made, with impedance of about 1 MOhm at 1 kHz. Either one or two microelectrodes were inserted into the cortex using a multi-electrode microdrive (NAN instruments). Neuronal signals were collected at 20 kHz, amplified using conventional differential amplifiers (CED 1902 Cambridge Electronics Design) and band-pass filtered (cut-off frequencies: high 300 Hz, low 5 kHz). We used a Schmitt-trigger to digitize the analog neuronal signal online into a computer-compatible TTL signal. We did not use the Schmitt-trigger to separate simultaneous recordings from multiple neurons, in which case we searched for another recording from only a single neuron, or we stored the data in analog form for offline separation by dedicated custom-written software in MATLAB. In addition to their anatomical location, putative dopamine neurons were identified following established criteria of low baseline activity rate (typically less than 8 impulses per second), strong phasic response to unexpected rewards (manually or automatically delivered at unexpected times) and elongated waveform of the action potentials (typically longer than 2 ms).

An infrared eye tracking system monitored eye position (ETL200; ISCAN), with temperature check on an experimenter’s hand at the approximate position of the animal’s head.

### Behavioral tasks and recorded neurons

We recorded the neuronal responses in the choice task described earlier, as well as in a single-option task, in which only one of the two choice options were presented to the animal. The only difference between single-option and choice tasks was in the presence of one or two options at cue presentation time, while the rest of the trial was identical, with the monkey required to select one side through a joystick movement and receiving the corresponding reward. The single-option task allowed for the investigation of basic processing aspects of value coding, avoiding potential confounds due to neuronal signals related to decisions or to combinations of the two options’ values. On the other hand, the choice task represented a situation more directly related to economic values during decision making, with the possibility of clearly identifying chosen value signals. Moreover, the choice task allowed us to simultaneously collect behavioral and neuronal subjective value measures, crucial for our direct comparison between the two quantities.

When a single neuron was identified and isolated, we recorded its activity during the single-option task with a limited set of reward parameters (safe option B: 0.15 ml or 0.35 ml, probability 1.0; gamble: magnitude 0.5 ml, probability 0.3 or 0.7). This was used to assess stability of the neuronal signal, neuron isolation, and to confirm typical dopamine reward responses. Then we ran the full set of reward parameters on these dopamine neurons. These parameters included 7 – 20 trial classes (including at least 3 safe options and 4 gambles) intermingled and equally presented for at least 8 repetitions (12 replications on average). In Monkey A, we typically recorded both single-option and choice trials intermingled, while in monkey B, due to shorter sessions, we mostly collected either single-option or choice trials. This resulted in the collection of activity from 172 dopamine neurons from two animals (N = 59 in monkey A, N = 113 in monkey B), with most monkey A’s neurons collected in both tasks (56 and 59 neurons in single-option and choice task respectively, for a total of 59 distinct neurons) and mostly separate groups of monkey B’s neurons (59 and 66 neurons in single-option and choice task respectively, for a total of 113 distinct neurons). We interrupted the recording session when a neuron’s waveform became too small or was lost during the recording. We included neurons that were well-isolated for at least 4 repetitions. When offline-sorting the signals, other neurons not observed during recordings were added to the database if their waveform was clearly different from that of other simultaneously isolated neurons and well separated from the background noise signal.

For a subset of dopamine neurons (N = 32 in Monkey A; N = 37 in monkey B), we offered the monkey choices between options which also included two- and three-outcome gambles. The specific gambles were defined based on the independence axiom of EUT (see below). This task was used to test the prediction of independence axiom violation due to the nonlinear probability weighting.

### Neuronal task relationships

Separately for single-option and choice tasks, we selected neurons with significant responses to both reward magnitude and probability at cue presentation. To identify this basic reward magnitude and probability modulation, we selected neurons for their cue responses varying with both reward magnitude and reward probability, as defined by significance of the multiple linear regression:

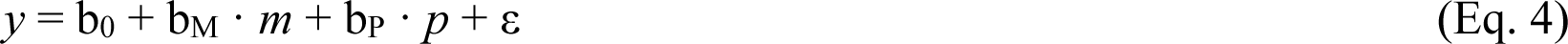

with *y* as mean activity rate (imp/s) calculated 100 to 300 ms after cue onset, b_0_ as offset, b_M_ and b_P_ as slopes for the regressors of magnitude (*m*, in ml) and probability (*p* ∈ [0,1]), respectively, ε as error term. In the choice task, m and p were those of the chosen option. This procedure identified dopamine neurons with both slope parameters b_M_ and b_P_ significantly different from 0 (t test, P < 0.05), which represented our data base for all subsequent data analyses.

### Elicitation of neuronal indifference points

Mirroring the behavioral utility measure, we defined a neuronal subjective economic value measure for each tested set of safe and gamble reward magnitudes. The neuronal indifference point (IPn) was defined as the gamble probability for which the neuronal response to the safe option matched the response to the gamble option. The neuronal indifference point (IPn) was defined as the gamble probability for which the neuronal response to the safe option matched the response to the gamble option (m = 0.5 ml). The IPn was computed as the intersection between two lines: the regression line of the neuronal activity for different gamble probabilities and the line representing the mean response to the safe option. A bootstrap procedure (see below) was used to determine the 95% confidence interval of the estimated IPn, in order to compare the neuronal measure with the equivalent behavioral measure (IPb).

### Nonlinear fit of neuronal responses to reward magnitude and probability

To characterize the neuronal responses to different levels of reward amount and probability separately, we fitted cubic spline functions to the neuronal population responses. This type of functions was selected to avoid arbitrary parametric assumptions which may bias fit results by forcing the curves to pre-defined shapes.

Using the *fit* Matlab function (method: *SmoothingSpline*; *SmoothingParam: 0.99996*) on the response rates we obtained cubic spline curves that approximated the average activities in response to either safe reward magnitudes or gamble reward probabilities (with fixed gamble magnitude m = 0.5 ml). We first fitted the responses to probabilities, and used the minimum and maximum spline fitted values as extreme points for the subsequent fit on the reward magnitudes. This was done to maximize the number of data points, and was in particular useful for magnitude domain for which the data resolution was lower than that of the probability domain. This procedure was justified by the fact that, by definition, the extreme points coincided between the magnitude and probability domains, as they represented either no reward (m = 0 ml or p = 0; leftmost point), or a safe option of 0.5 ml (rightmost point). The fit parameter was used to obtain smooth fits which closely followed the data points, without imposing any further condition of monotonicity on the fitted functions.

As the presented options varied across sessions due to slight adjustments in the task composition, for this data analysis we only included neurons that were recorded during the presentation of the same set of choice options. Despite reducing the number of neurons for this analysis, this approach avoided the introduction of additional sources of data variability.

This procedure was applied to the population activity (Figure 5 and 6) as well as to the individual neurons (Figure 8).

### Using the independence axiom for testing neuronal value code predictions

The fourth axiom of EUT, the independence axiom, defines how reward magnitudes and probabilities are combined into a scalar subjective value, expected utility (EU). According to EUT, the EU of a gamble is the sum of all outcomes’ utilities, weighted by their respective probabilities:

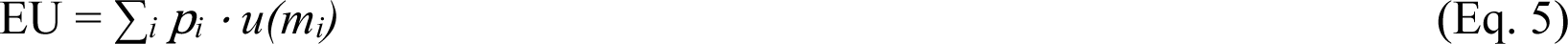

where *i* is the outcome index (from 1 to the number of outcomes), *u* the utility function (representing the subjective transformation of objective reward magnitudes), and *p* the objective reward probability.

Compliance with the axiom is required for this value formulation to represent choices. According to the axiom, preferences between two gambles A and B should not change after adding a common probability of the same outcome to both options. This can be empirically tested by defining two gambles C and D as modified versions of the original A and B gambles. A subject preferring A to B (or B to A), should also prefer C to D (or D to C). Any other combination of preferences would violate the axiom. Formally the axiom states that

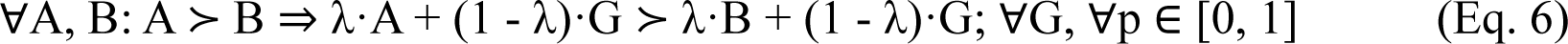

where A, B represent the original gambles; G represents a gamble (or a safe outcome) that commonly extends both original choice options, weighted by λ (lambda), which represents the probability of the original A and B gambles within the new C and D gambles. In our design, gamble A was a safe option as defined in the original test by Allais (Allais, 1953). We assessed IA compliance in two commonly used tests: the common consequence (CC) test and the common ratio (CR) test. The CC test consisted of adding (or subtracting) the same specific probability of an outcome (’common consequence’) commonly to both options, exactly as prescribed by the axiom. The CR test consisted of multiplying the same ’common ratio’ with the probabilities of all non-zero outcomes commonly for both gambles A and B. Changes in preferences between the {A, B} and {C, D} option pairs in both CC and CR tests correspond to axiom violations.

When the axiom is violated, the EU formulation does not describe choices. Following empirical evidence for axiom violation, non-EU theories have been developed (Starmer, 2000). Prominently, prospect theory (PT) (Kahneman & Tversky, 1979) modified the value formulation to allow for nonlinear probability weighting, resulting in the following ‘prospect value’ (i.e., gamble value) equation:

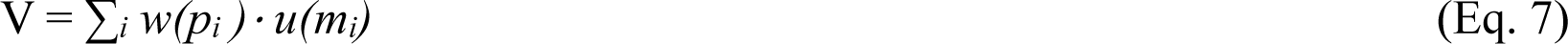

where *w* is the probability weighting function, i.e., a subjective transformation of the objective probabilities. For gambles with more than one nonzero outcome, the prospect value was formulated in cumulative PT (Tversky & Kahneman, 1992) as

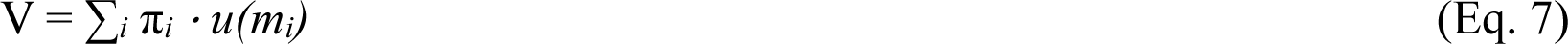

where π is a combination of probability weights which depends on the ranking, in terms of magnitudes, of the gamble outcomes. This component, which can be expressed as

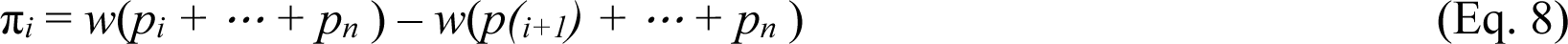

was introduced to extend the original PT formulation to more complex choice options without incurring in basic problems such as the violation of stochastic dominance. We used this cumulative PT approach when computing the neuronal indifference curves within the Machina triangle (see below).

Regardless of the specific formulation for the value function, the contribution of the independence axiom to our study lies in the prediction of specific patterns of choices (axiom violations) from the assumption of nonlinear probability weighting.

### Neuronal indifference curves

To compute the neuronal indifference curves (ICs), we first computed a neural value quantity for each point (i.e., gamble) in the magnitude probability space. We used a resolution of 0.01 for both magnitude (0 to 0.5 ml) and probability (0 to 1) dimensions. Assuming the original PT value formulation (Eq. 7), we computed values from the neuronal utility and weighted probability functions. We assumed that the nonlinear fit of neuronal responses to magnitudes and probabilities represented the utility and weighted probability function, respectively. We then combined these quantities for each gamble, obtaining a value map across the tested magnitude-probability space. Finally, we computed the indifference curves as points with the same value (Matlab function: *contour3*).

We compared neuronal and behavioral indifference curves using the mean squared error as metric, in the probability direction. The error was computed between each measured IPb and the neuronal IC point corresponding to the same magnitude level. The mean squared error was computed across all tested IPs within the indifference map calculated for each animal separately.

The same procedure was repeated (Figure 6) assuming the linear coding of probabilities (Eq. 5) and also assuming an additive combination of neuronal magnitude and probability responses (V = ∑_i_ *w(p_i_) + u(m_i_)*). Results following these different assumptions were compared in terms of the distribution of mean squared errors, computed across bootstrapped resamples (see below for details on the bootstrap procedure). Using the same approach, we also compared the mean squared error between neuronal ICs and the other animal’s behavioral ICs.

Neuronal ICs were also computed within the Machina triangle, assuming a cumulative PT representation of weighted probabilities (Eq. 7).

### Statistical analysis

All statistical tests (t-test, Kolmogorov-Smirnov test, correlation (Pearson), binomial test) were considered significant at the α = 0.05 level. We used a bootstrap procedure to compute the error areas associated with the following measures: behavioral indifference curves (Figure 1D); neuronal population responses to magnitude and probability (Figure 2C, D; Figure 5 A, B); neuronal indifference points and fitted curves (Figure 3C, D); indifference curves computed as combination of neuronal utility and probability weighting functions (Figure 5C, D). For the bootstrap procedure on the behavioral data, we resampled with replacement a fixed number of trials (N = 10) for each choice pair, to simulate between-session variability. To perform the bootstrap procedure on the neuronal data, we resampled with replacement the average activity of individual neurons and then computed the above measures using the resampled data. The procedure was repeated 1,000 times and the standard deviations and 95% CIs (Matlab function: *prctile*) from the resulting measures were calculated. For a clean graphical representation, only a random subset of 100 resamples was plotted.

